# Integrative Multi-Omics Analysis Identifies Nuclear Factor I as a Key Driver of Dysregulated Purine Metabolism in DIPG

**DOI:** 10.1101/2025.05.21.655327

**Authors:** Ian Mersich, Sunny Congrove, Matthew Horchar, Roman Caceres, Ranjithmenon Muraleedharan, Janki Desai, Pankaj Desai, Larry Sallans, Julie A. Reisz, Abby Grier, Matthew R. Hass, Omer Donmez, Cailing Yin, Matthew T. Weirauch, Leah Kottyan, Charles B. Stevenson, Claire Sun, Peter de Blank, Natasha Pillay-Smiley, Trent R. Hummel, Nagarajan Elumalai, Ali Tavassoli, Ron Firestein, Timothy N. Phoenix, Angelo D’Alessandro, Biplab Dasgupta

**Author notes:** **Corresponding Author and Lead Contact Contact: Biplab Dasgupta PhD**, MBA Professor of Pediatrics-Neuro-oncology Emory University School of Medicine 1750 Haygood Drive Atlanta, GA, 30322. Equal contribution (Matthew Horchar, Roman Caceres, Ranjithmenon Muraleedharan). Department of Chemistry and Biochemistry, University of Notre Dame, IN, USA. Center for Development of Therapeutics (CDoT), Broad Institute of MIT and Harvard, Cambridge, MA, USA.

## Abstract

Diffuse intrinsic pontine glioma (DIPG) is a devastating brainstem cancer in children, with a median survival of under one year and limited treatment options. Over 80% of DIPGs possess a H3K27M mutation. To identify metabolic vulnerabilities linked to this mutation, we utilized a multi-omics approach in H3K27M-expressing cells, patient-derived cell lines, and mouse models. We show that by reprogramming chromatin landscape the mutation aberrantly induces NFI transcriptional activity, leading to misregulated purine metabolism. The mutation amplifies purine biosynthesis and degradation via the enzymes ATIC and PNP, respectively. Unregulated purine degradation relieves the negative feedback of purines on their own synthesis allowing continuous synthesis, use and degradation making DIPGs reliant on purine biosynthesis. Targeting ATIC reduced tumor progression and improved survival in mice. We propose ATIC as a potential novel target in DIPG.

## INTRODUCTION

Diffuse intrinsic pontine glioma (DIPG) is one of the most lethal pediatric malignancies. Due to the anatomical location, surgical resection is not possible. Moreover, radiation and conventional chemotherapies provide little to no extension in life leading to a poor median overall survival of just 11 months^1^. Throughout the last decade, research has greatly improved our understanding of the molecular landscape of these tumors including the identification and classification of the oncogenic K27M mutation in histone 3 genes (H3K27M) in most patients ^2,3^. H3K27M directly alters the epigenetic landscape in cells through the dampening of PRC2 repression and global reduction of H3K27me3 resulting in aberrant gene expression, driving oncogenic reprograming^4–6^. Despite advancements in understanding the pathobiology of DIPG, and the identification of novel targeted treatment strategies^7–10^, clinical trials have yet to translate these findings into a significant increase in progression free survival or overall survival in patients^11^. The poor prognosis underscores an urgent, yet unmet, need to further understand the molecular intricacies of DIPG pathobiology and mechanisms of therapeutic resistance.

Metabolites serve as proxies to the state of biochemical pathways in a tissue, and reprogramming of metabolic pathways to support rapid growth, proliferation and survival is a signature of cancer^12,13^. Initiated by the pioneering work using anti-folates in treating childhood ALL^14^, drugs that target metabolic pathways have successfully been used to treat malignancies including L-asparaginase, routinely used for treating ALL^15^, and anti-metabolites targeting nucleotide biosynthesis^16,17^; however, many promising preclinical studies have failed in the clinic due to a limited therapeutic window in pathways essential to normal healthy tissue such as glycolysis^18,19^, and glutamine metabolism^20,21^. Given the unique metabolic demands of the central nervous system and the intricate genetic landscape of DIPGs, it is plausible that these tumors harbor distinct metabolic reprogramming events pivotal to their aggressive nature.

Research on altered metabolism in DIPG is limited; however, it is becoming evident that H3K27M mutations play a significant role in driving metabolic changes. Expression of the mutation alone in neural stem cells enhances glycolysis, glutaminolysis, and TCA cycle metabolism, elevating levels of alpha-ketoglutarate shown to be necessary to maintain low global H3K27me3 levels^22^, a defining feature in H3K27M gliomas. This research aligns with previous clinical observations. Using in vivo proton magnetic resonance spectroscopy (1H MRS), DIPG patients exhibited elevated citrate levels in tumor tissue compared to normal tissue and when contrasted with other pediatric brain tumors^23–25^. Glycolytic shunt into the pentose phosphate pathway has previously been reported as a direct consequence of the H3K27M mutation^26^. Phosphoribosyl diphosphate (PRPP), a product of the pentose phosphate pathway, is essential for nucleotide biosynthesis and DIPGs have been reported to be dependent on de novo pyrimidine biosynthesis^27^.

Given that knowledge about metabolic alterations in DIPG is limited, we set out to identify metabolic dependencies inherent to the H3K27M mutation. To gain a more comprehensive view of altered metabolism driven by the mutation and address the physiological relevance, we leveraged a multi-omics analysis of H3K27M-expressing cells, including a panel of DIPG cell lines and tumor tissue in DIPG mouse models. Our comprehensive profiling enriches our understanding of DIPG pathogenesis, connecting epigenetic abnormalities stemming from the mutation to gene regulatory networks, which in turn influence metabolic alterations and unveil therapeutic vulnerabilities.

## RESULTS

### Purine Metabolism Is Consistently Dysregulated Across Human, Murine, and PDX Models of DIPG

Untargeted metabolomics was performed on six genetically characterized patient-derived DIPG lines and one normal human neural stem cell (NSC) line to identify metabolic alterations and therapeutic targets. The DIPG lines segregated into two distinct clusters, each notably different from NSCs (Figure 1A). Enrichment analysis of the most consistently altered metabolites across all DIPG lines identified several pathways of interest (Figure 1B, Supplemental Table S1). In parallel, RNA sequencing of these lines was conducted to identify differentially expressed genes (DEGs) associated with metabolic alterations.

**Figure 1.**
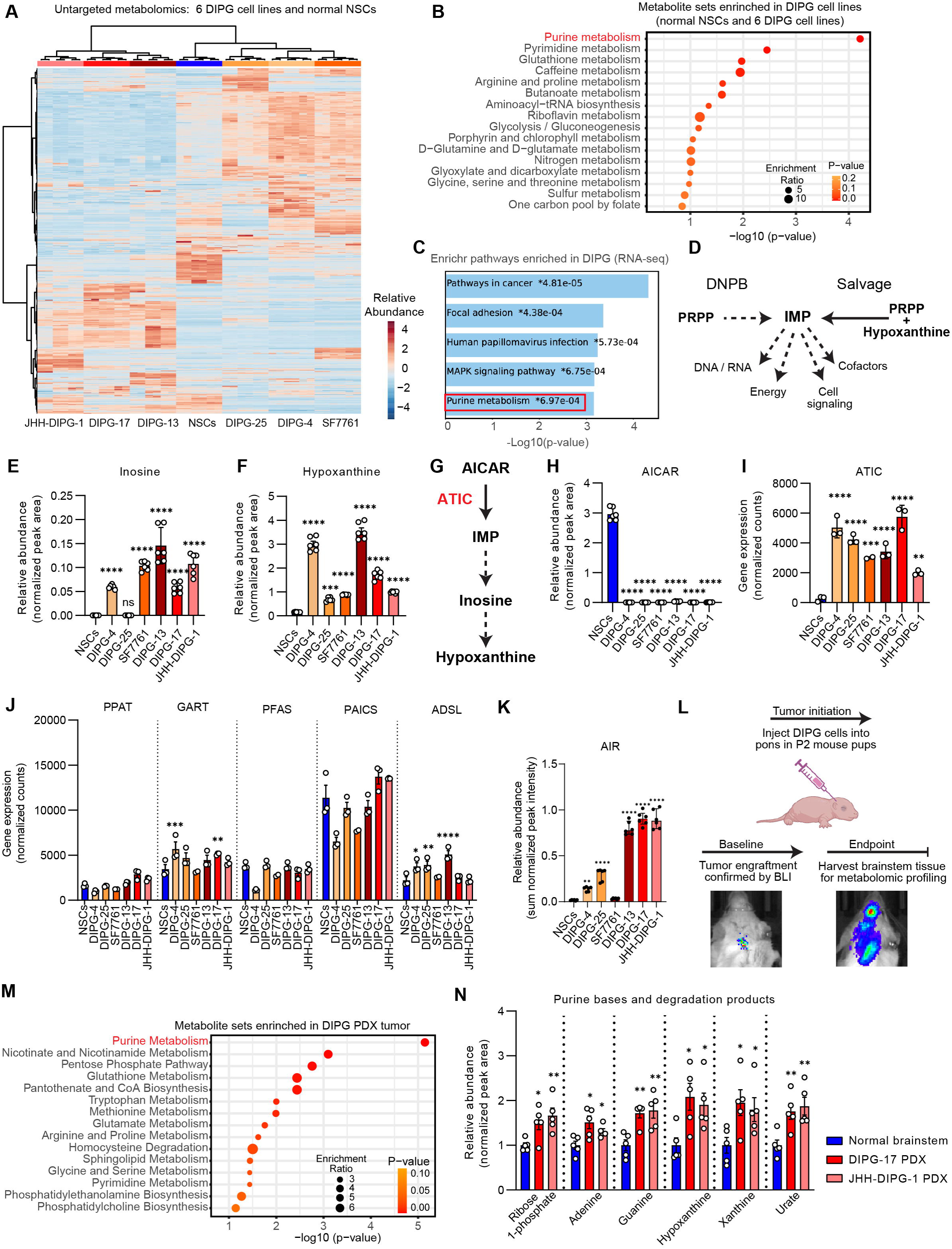
**Purine metabolism is consistently dysregulated across human, murine, and PDX models of DIPG** (A) Heatmap of untargeted metabolomics analysis in 6 H3K27M DIPG cell lines and 1 human NSC cell line (n=6) (B) Pathway enrichment analysis of differentially regulated metabolites in DIPG vs NSCs (P < 0.05; fold change ≤ 0.5 or ≥ 2). (C) Enrichr gene set enrichment plot for pathways in KEGG_2021_Human database from DEGs in 6 DIPG cell lines vs human NSCs (n=3 each). (D) Schematic depicting the cellular functions of purines and production of the AMP/GMP precursor IMP from PRPP through de novo biosynthesis or recycled through the salvage pathway. (E and F) Relative abundance (sum normalized peak area) of inosine and hypoxanthine in DIPG vs NSCs from untargeted metabolomics. (G) Schematic depicting ATIC as a rate-limiting step through the conversion of AICAR to IMP and downstream metabolites. (H) Relative abundance (sum normalized peak area) of ATIC substrate AICAR from untargeted metabolomics comparing each DIPG cell line individually with NSCs (n=6) (I) Relative gene expression (normalized counts) of DNPB enzyme ATIC from RNA-sequencing analysis (n=3). (J) Relative expression of other DNPB genes across DIPG and NSCs (n=3). (K) Relative abundance of AIR metabolite in DIPG vs NSCs (n=4-6). (L) Schematic workflow of DIPG-PDX generation and in vivo metabolomics profiling. (M) Enrichment of metabolites in DIPG-PDX tumors vs normal brainstem. (N) Relative abundance (sum normalized peak area scaled to control) of purine bases and degradation products in DIPG PDX tumors. Tumors derived from two separate DIPG cell lines individually compared to normal mouse brains (n=5). Panels E, F, H, I, J, and K plotted as mean ± SD, analyzed by one-way ANOVA with Dunnett’s correction. Panels M and N plotted as mean ± SEM; comparisons analyzed by unpaired two-tailed T test. (* P < 0.05; ** P < 0.01; *** P < 0.001; **** P < 0.0001).

Gene set enrichment analysis (GSEA) and a joint pathway analysis integrating metabolites and DEGs revealed that differences between the two DIPG clusters were primarily related to lipid metabolism (Figures S1A and S1B). Separately, we performed an enrichment analysis on all DEGs in DIPGs compared to NSCs to identify pathways enriched in DIPGs (Supplemental Table S2-4). EnrichR enrichment analysis identified gene sets expected to be enriched in DIPGs related to PDGFRA and PRC2 (Figure S1C). Among these, purine metabolism was one of the most significantly enriched gene sets related to metabolic pathways (Figure 1C). A KEGG Pathview diagram provided a broad overview of purine metabolism, demonstrating extensive alterations of genes and metabolites in DIPG cells relative to NSCs (Supplemental File 1).

Purine nucleotides are synthesized via de novo purine biosynthesis (DNPB) or salvaged from purine bases (Figure 1D). Inosine and hypoxanthine levels were significantly elevated in DIPGs compared to NSCs (Figures 1E and 1F). Most importantly, we identified a critical rate-limiting step in DNPB that may be dysregulated in DIPGs. Specifically, the metabolite AICAR used as a substrate by the enzyme 5-aminoimidazole-4-carboxamide ribonucleotide formyltransferase/IMP cyclohydrolase (ATIC) at the last step of DNPB to produce inosine monophosphate (IMP) (Figure 1G) was downregulated in DIPGs (Figure 1H). Markedly lower AICAR levels (Figure 1H) and elevated ATIC expression in DIPGs (Figure 1I) suggest rapid utilization of this metabolite for downstream purine biosynthesis. Although not all DNPB genes were consistently upregulated across all DIPG cell lines (Figure 1J), the levels of the DNPB intermediate AIR was elevated in DIPG cells compared to NSCs (Figure 1K). Collectively, these findings suggest that high ATIC expression contributes to enhanced purine biosynthesis and increased production of downstream purine metabolites in DIPG.

Dysregulated purine metabolism was further examined using cell lines derived from a genetically engineered mouse model of DIPG generated via brainstem-targeted in utero electroporation (IUE) of plasmids encoding mutant PDGFRA (PDGFRA D842V), dominant negative P53 (DNP53), and mutant histone H3 (H3.3K27M). Untargeted metabolomics confirmed that this defined oncogenic combination led to enrichment of purine metabolites in the resulting mouse lines (Figure S1D). Integration of these metabolomics data with previously acquired RNA-seq profiles confirmed purine metabolism as one of the most significantly altered pathways at both the gene and metabolite levels (Figure S1E, Supplemental Table S5). Unlike human DIPG cell lines, hypoxanthine levels in the murine DIPG cell lines were comparable to the normal mouse NSCs; however, other purine degradation products—including inosine, 5′-hydroxyisourate, and allantoin—were consistently elevated in IUE tumor cells (Figure S1F). These results suggest that although individual metabolite levels may fluctuate due to metabolic flux, overall dysregulation of the purine pathway is preserved across models.

Next, we performed in vivo analysis was to determine whether purine metabolism is also altered in tumor tissue. Two DIPG patient-derived xenograft (PDX) models were generated by orthotopic implantation of luciferase-labeled JHH-DIPG-1 and SU-DIPG-17 cells into the brainstem of postnatal day two mice (Figure 1L). Untargeted metabolomics of these tumors and normal brainstem tissue revealed that the DIPG-PDX samples were more similar to each other than to normal brain, as shown by partial least squares discriminant analysis (PLS-DA) (Figure S1G). Consistent with in vitro findings, DIPG tumors displayed broad metabolic changes in central carbon pathways, including dysregulation of purine metabolism (Figure 1M). Specifically, purine bases and degradation products were elevated in DIPG tumors (Figure 1N), while nucleoside and nucleotide levels remained comparable to those in normal brain tissue (Figure S1H). The consistent absence of elevated purine nucleotide levels, despite the presence of purine bases and degradation products across all three models, suggests that excess nucleotides are rapidly degraded to prevent feedback inhibition of DNPB.

Potential conversion of purine nucleotides into cyclic second messengers was also considered. While cAMP and cGMP was not detected in DIPG cell lines by untargeted metabolomics, and levels varied across DIPG-PDX tumors and normal brain samples, transcript levels for multiple isoforms of guanylate cyclases (GUCY), adenylate cyclases (ADCY), and phosphodiesterases (PDE) were significantly elevated in DIPG lines compared to NSCs (Figure S1I, See also Supplemental Table S2 and S3), supporting the possibility that a subset of purines is diverted into second messenger pools in DIPG.

Additional dysregulated pathways were identified across all three DIPG models. Metabolites from the methionine cycle (cysteine and choline), which is functionally connected to DNPB via the folate cycle, and ornithine, a key substrate for polyamine biosynthesis, were consistently elevated in DIPG cell lines (Figures S1J-L), PDX tumors (Figures S1M-O), and IUE-derived tumor cells (Figures S1P-R). These findings highlight strong cross-model congruence in auxiliary metabolic alterations associated with DIPG.

### Altered purine metabolism in DIPG is driven by unregulated purine biosynthesis and degradation pathways

Given the high expression of ATIC, the corresponding depletion of its substrate AICAR, and elevated levels of intermediate purines and degradation products—including hypoxanthine and inosine, a nucleoside positioned between de novo biosynthesis and salvage—we hypothesized that both the de novo and salvage pathways contribute to the accumulation of purine metabolites in DIPG. Inosine may act as a purine reservoir, supporting unregulated DNPB activity, particularly in the context of known nucleotide-mediated feedback inhibition at multiple steps of the purine biosynthesis pathway.

To examine the relative contributions of DNPB and salvage pathways to the purine pool in DIPG, we next performed stable isotope tracing experiments. Three separate tracer experiments were conducted using [¹³C_₆_]glucose and [¹[N]serine to monitor DNPB activity, and [¹[N_₄_]hypoxanthine to assess salvage contributions (Figure 2A). As glycine serves as a nitrogen donor in DNPB, the incorporation of labeled nitrogen from serine into glycine and subsequently into downstream purines—including inosine were measured. Tracer incorporation in glycine and inosine was elevated in DIPGs while that to AICAR was decreased (Figures 2B and 2C), consistent with our in vitro untargeted metabolomics data. Notably, DIPG lines exhibited markedly increased levels of labeled urate (Figure 2D), a purine degradation product which is also in line with our in vitro tumor metabolomics data (see Figure 1N). Tracing with [¹[N_₄_]hypoxanthine revealed that purine salvage is functional and also contributes to purine pools through the incorporation into inosine and degradation to urate (Figures 2E–G). In line with established nucleotide-mediated feedback inhibition, the abundance of most labeled adenylates and guanylates remained relatively unchanged between DIPG lines and NSCs (Figures S2A–D). An additional [¹³C_₆_]glucose tracing experiment confirmed unrestricted carbon flow from glucose into inosine through the DNPB pathway (Figure S2E). Together, these experiments show active contribution of both DNPB and salvage pathways to purine synthesis and degradation of excess purines.

**Figure 2.**
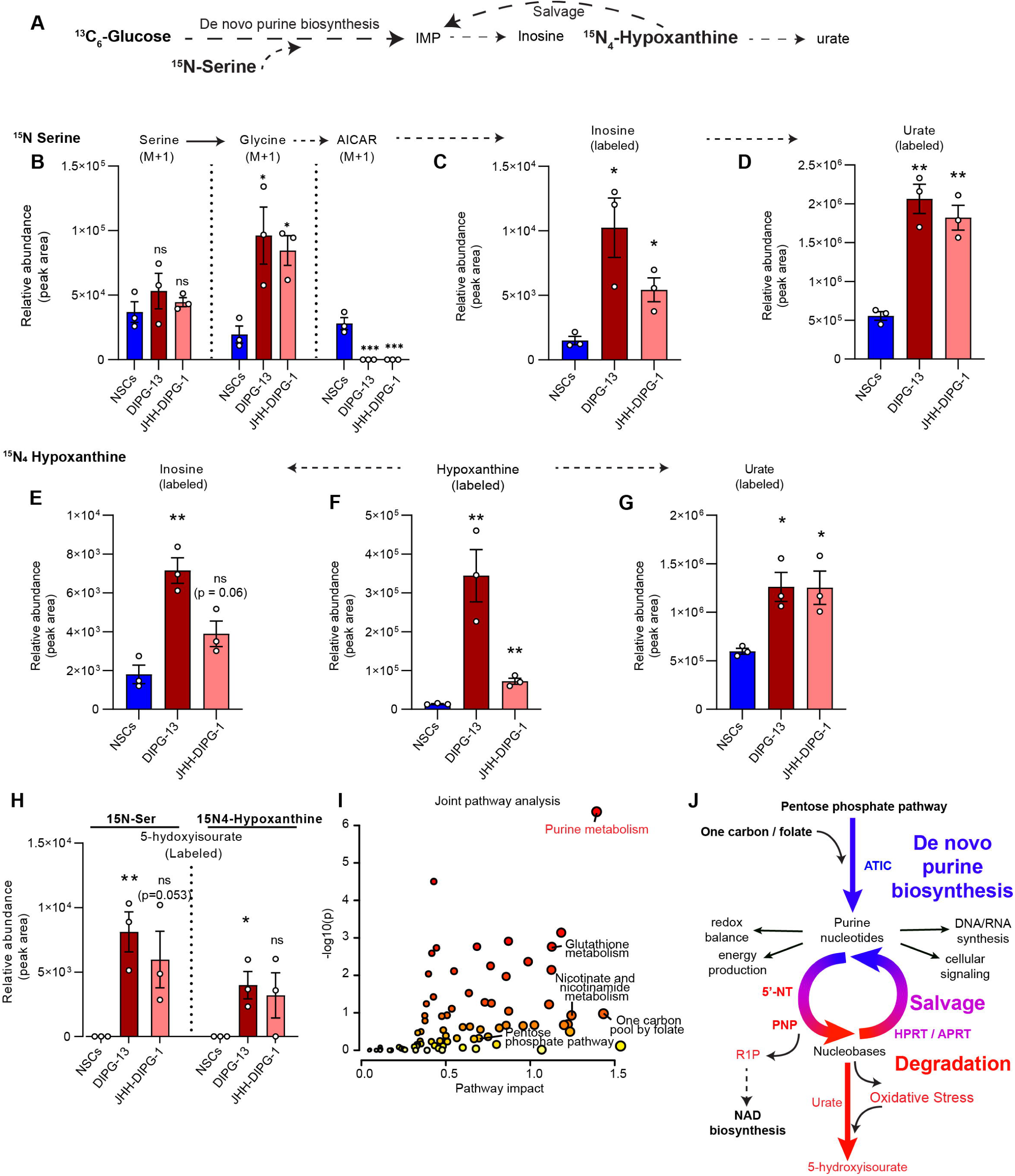
**Altered purine metabolism in DIPG is driven by unregulated purine biosynthesis and degradation pathways.** (A) Schematic of isotope tracing workflow using [13C6]glucose, [15N]serine, and [15N4]hypoxanthine to distinguish contributions from DNPB and salvage pathways. (B) Relative abundance (peak area) of 15N-labeled (M+1) serine, glycine and AICAR from NSCs and 2 DIPG cell lines (n=3) incubated with 15N-serine (See also Figure S2). (C-D) Relative abundance (peak area) of labeled fractions for the purine intermediate inosine and the purine degradation product urate from panel B. (E-G) Same as panels B-D from cells incubated with 15N4-hypoxanthine. (H) Relative abundance (peak area) of labeled 5-hydroxyisourate from both tracing experiments. (I) Enriched pathways from Metaboanalyst joint pathway analysis, combining differentially expressed genes and differentially regulated metabolites (see also Supplemental table S1 and S2) (J) Schematic highlighting key genes facilitating purine biosynthesis (blue) salvage (purple) and degradation (red) and cognate metabolites that are consistently high in DIPGs relative to NSCs. Pathways closely connected to purine metabolism and enriched in DIPGs (from Figure 2I) indicated in bold. The oxidation of hypoxanthine to xanthine, and then to urate generates oxidative stress, and this likely creates a feedforward loop for the oxidation of urate to 5-hydroxyisourate. Panels B-H plotted as mean +/- SEM. Each DIPG line was compared to NSCs, analyzed by unpaired, two-tailed T test (ns = not significant; * p < 0.05; ** p < 0.01; *** p < 0.001)

We noticed that the levels of the urate oxidation product 5-hydroxyisourate were significantly elevated in DIPG cells relative to NSCs, which was consistent across multiple experiments (Figures S2F–H). These findings indicate that increased purine degradation, evidenced by high levels of urate and 5-hydroxyisourate in our tracing experiments (Figure 2H), relieves feedback inhibition and permits unrestricted DNPB activity. Consistently, multiple genes involved in purine degradation and salvage—including purine nucleoside phosphorylase (PNP), hypoxanthine-guanine phosphoribosyltransferase (HPRT), adenine phosphoribosyltransferase (APRT), guanosine monophosphate reductase (GMPR), and several nucleotidases—were highly expressed in DIPG lines, indicating coordinated regulation of purine metabolism at both the transcript and metabolite levels (Figures S2I–L, Supplemental Table S2).

Due to the absence of a functional uricase enzyme in humans, purine catabolism proceeds through urate and 5-hydroxyisourate, driven by oxidative stress and generating reactive oxygen species(ROS)^34–36^. Indeed, metabolomic profiling of DIPG lines consistently revealed markers of oxidative stress, including a decreased reduced-to-oxidized glutathione ratio (GSH/GSSG) and elevated dehydroascorbate levels (Figures S2M–P). Transcriptomic data further confirmed upregulation of ROS-associated genes and antioxidant response genes in H3K27M DIPG patient tumors compared to H3 wild-type high-grade glioma (HGG) and low-grade glioma (LGG), and in DIPG lines compared to NSCs (Figures S2Q–R). In support of these findings, DIPG lines and NSCs ectopically expressing the H3K27M mutation exhibited higher levels of total and mitochondrial ROS relative to H3WT-expressing NSCs (Figures S2S–T). These data collectively indicate that elevated oxidative stress and enhanced ROS buffering capacity are hallmark features of DIPG, supporting the spontaneous oxidation reactions required for purine degradation.

A joint pathway analysis, integrating differentially expressed genes and metabolites from RNA-seq and untargeted metabolomics, was used to investigate how dysregulated purine metabolism connects to broader metabolic alterations (Figure 2I). This analysis identified enrichment of DNPB-supporting pathways such as the pentose phosphate pathway and folate metabolism (Figure 2J). Additionally, it revealed functional connections to pathways fueled by purine degradation, including the generation of ribose-1-phosphate (R1P) by PNP, which can be utilized for NAD biosynthesis—potentially compensating for increased oxidative burden in DIPG (Figure 2J).

Collectively, these in vitro and in vivo findings from untargeted metabolomics and isotope tracing establish that dysregulated purine metabolism in DIPG is characterized by unchecked DNPB, rapid AICAR turnover by the rate-limiting enzyme ATIC, recycling of intermediates through salvage enzymes, accelerated purine degradation to overcome nucleotide-mediated feedback inhibition, and elevated oxidative stress (Figure 2J).

### H3K27M mutation drives dysregulated purine metabolism through upregulation of ATIC and purine pathway genes

The transcriptomic and metabolomic analyses across multiple in vitro and in vivo models revealed consistent upregulation of DNPB and purine degradation in DIPG. Analysis of patient RNA-seq data from the PNOC003 cohort (PedcBioPortal) showed significantly higher expression of ATIC in DIPG tumors relative to normal brain tissue from GTEx (Figure 3A). Three other DNPB genes GART, ADSL and PAICS were also upregulated (Figure 3A). Similarly, across pediatric brain tumors in the OpenPBTA cohort, patients with the H3K27M mutation showed elevated expression of ATIC (Figure 3B), and other DNPB genes (S3A–E) compared to H3 wild-type tumors. Consistent with the enhanced purine degradation phenotype observed in our metabolomics data, expression of the purine degradation enzyme PNP was also elevated in H3K27M tumors (Figure S3F).

**Figure 3.**
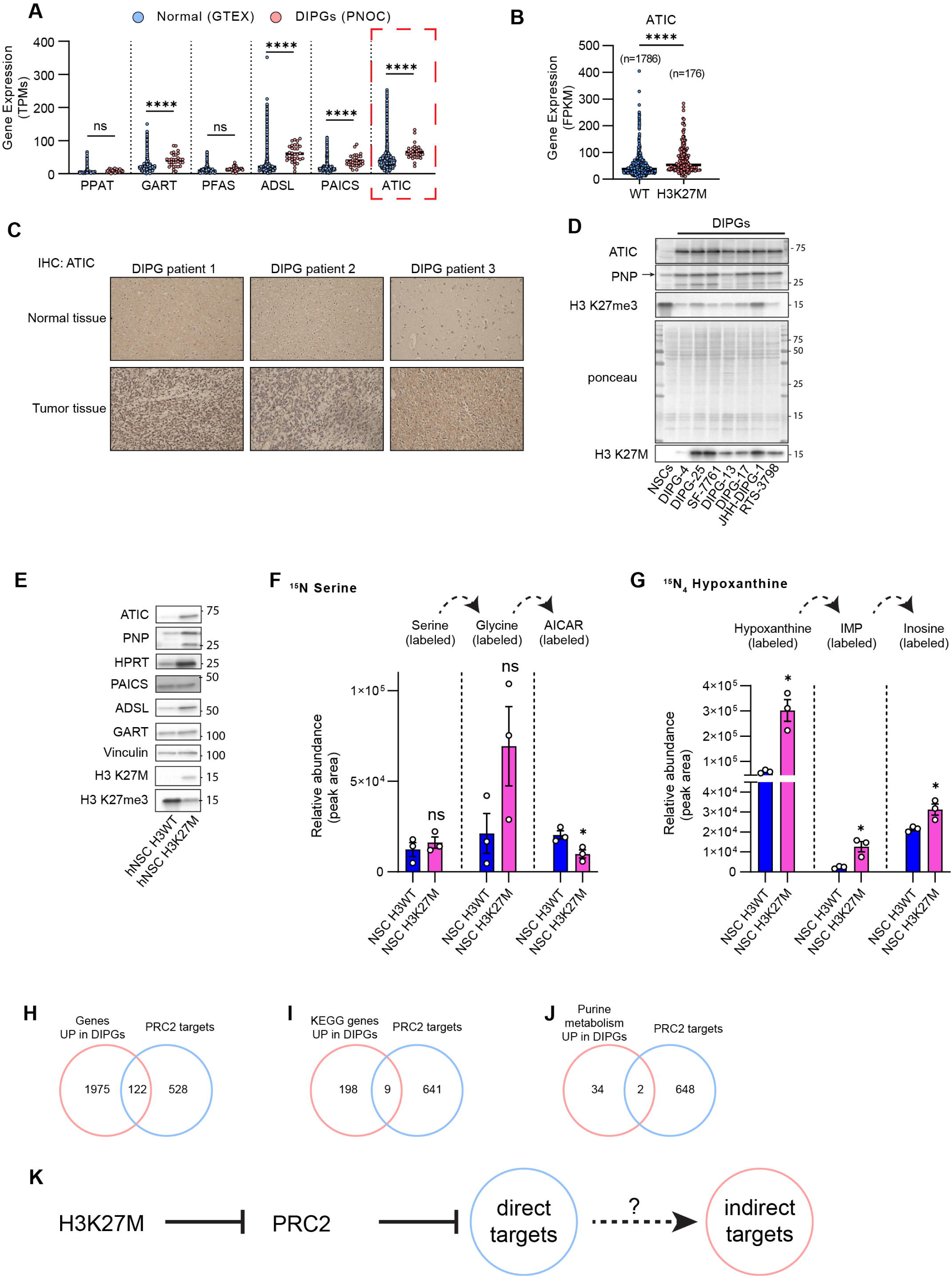
**H3K27M mutation drives dysregulated purine metabolism through upregulation of ATIC and purine pathway genes** (A) Relative gene expression (TPMs) for DNPB genes including ATIC in normal brain (GTEx, n=17382) and DIPG (PNOC003, n=32). (B) ATIC expression across pediatric brain tumors with and without H3K27M mutation. (C) Representative immunohistochemistry (IHC) showing ATIC protein expression in DIPG patients. Tumor section (bottom) and matched normal tissue section from each (n=8). (D) Western blot comparing protein levels for ATIC and PNP in NSCs relative to DIPG cell lines, ponceau staining for total protein used due to varying levels of housekeeping genes across cell types. (E) Western blot comparing protein levels for multiple purine metabolism genes in H3WT NSCs relative to H3K27M NSCs. (F and G) Relative abundance (peak area) of 15N-labeled metabolites in NSCs expressing H3WT or H3K27M, (n=3) incubated with 15N-serine or 15N4-hypoxanthine (See also Figure S2). (H-J) Venn diagram of differentially expressed genes overlapping with known PRC2 targets among all DEGs (H), KEGG metabolic pathway genes (I), and purine pathway genes (J). (K) Proposed model where the H3K27M mutation alters gene regulation indirectly through transcription factor activation, including metabolic rewiring. Panels A-B plotted as scatter dot plot with line at median, analyzed by one-way ANOVA (A) and two-tailed T test with Welch’s correction (B). Panels F and G plotted as mean +/- SD, each mutant compared to H3WT NSCs analyzed by unpaired, two-tailed T test. (ns = not significant; * p < 0.05; ** p < 0.01; *** p < 0.001)

The clinical relevance of purine pathway activation in pediatric brain tumors was assessed and, survival outcomes were analyzed in relation to ATIC expression levels. High ATIC expression was associated with worse survival patients (Figure S3G). High ATIC expression is also associataed with shorter survival times in DIPG patients (Figure S3H); however, the effect was modest -likely due to higher overall expression of ATIC in DIPGs. These findings suggest that ATIC not only plays a functional role in purine metabolic dysregulation but may also serve as a biomarker of disease aggressiveness and a potential therapeutic target. Immunohistochemistry confirmed robust ATIC protein expression in human DIPG tumors (Figure 3D) and in orthotopic DIPG patient-derived xenograft (PDX) tumors (Figure S3I). In the IUE mouse model of DIPG, several DNPB and purine degradation genes were significantly upregulated in cell lines derived from tumor tissue relative to normal brainstem NSCs (Figure S3J and K), which was confirmed at the protein level by western blot (Figure S3L). Further, western blotting of DIPG lines and normal NSCs confirmed substantial upregulation of ATIC and PNP in all DIPG lines (Figure 3D). The H3K27M mutation, present in over 80% of DIPG cases, is a defining oncogenic event that initiates disease pathology. To test if the H3K27M mutation is sufficient to drive dysregulated purine metabolism, we made isogenic cell linesby stably expressing either the mutant or wild-type H3 histone in normal human NSCs via lentiviral transduction. NSCs expressing H3K27M demonstrated significant upregulation of ATIC, PNP, and the salvage enzyme HPRT (Figure 3E). Stable isotope tracing using [¹[N]serine and [¹[N_₄_]hypoxanthine was performed on these lines to functionally interrogate DNPB and salvage flux. Similar to DIPG cell lines, AICAR levels were lower in mutant NSCs, indicating increased consumption by ATIC (Figure 3F). Hypoxanthine tracing demonstrated an increase in the salvage pathway with an increase in labeled IMP, and degradation with an increase in labeled inosine (Figure 3G).

These results demonstrate that the H3K27M mutation alone is likely sufficient to induce purine metabolic rewiring in human NSCs. Whether altered metabolism in DIPG—specifically dysregulated purine metabolism—results from PRC2 inhibition by the H3K27M mutant histone was assessed by cross-referencing differentially expressed genes (DEGs) upregulated in DIPG with curated PRC2 target datasets from the Molecular Signatures Database (MSigDB). Only a small fraction of DEGs (6%, 122 out of 2,097) were predicted PRC2 targets (Figure 3H, Supplementary Table S6). Among DEGs involved in metabolism related pathways (KEGG), 4% (9/207) were predicted PRC2 targets (Figure 3I), and within the purine metabolism pathway, only 5% of upregulated genes (2/36) overlapped with known PRC2 targets (Figure 3J). These findings suggest that while PRC2 derepression may account for a subset of gene expression changes, the majority of metabolic gene upregulation in DIPG likely arises indirectly through secondary transcriptional network changes, wherein the H3K27M mutation enables aberrant activation of downstream signaling pathways or transcriptional programs (Figure 3K).

### NFI transcription factors mediate transcriptional activation of purine metabolism genes downstream of H3K27M mutation

The limited number of purine metabolism genes identified as direct PRC2 targets suggested an indirect mechanism of transcriptional activation caused by H3K27M mutations. From our RNA-seq data, the most significantly upregulated transcription factors in DIPGs were cross-referenced with transcription factor (TF) binding predictions for genes of interest from GeneHancer. Among these, the nuclear factor I (NFI) family of transcription factors emerged as top candidates for regulating ATIC. Importantly, the NFI TFs were identified as PRC2 targets in several curated gene sets and hence is expected to be aberrantly activated.

The connection between NFI-TFs and altered metabolism in DIPG was further investigated because they’ve been shown to be key regulators of glial and neuronal differentiation, critical for normal brain development[40,41], and are linked to altered metabolism[42]. RNA-seq data showed that all four NFI isoforms (NFIA, NFIB, NFIC, and NFIX) were significantly upregulated in DIPG cell lines compared to NSCs (Figure 4A), and in NSCs expressing the H3K27M mutant histone by qPCR (Figure S4A), suggesting a direct link between the mutation and NFI expression. This was also confirmed at the protein level by western blot analysis (Figure 4B).

**Figure 4.**
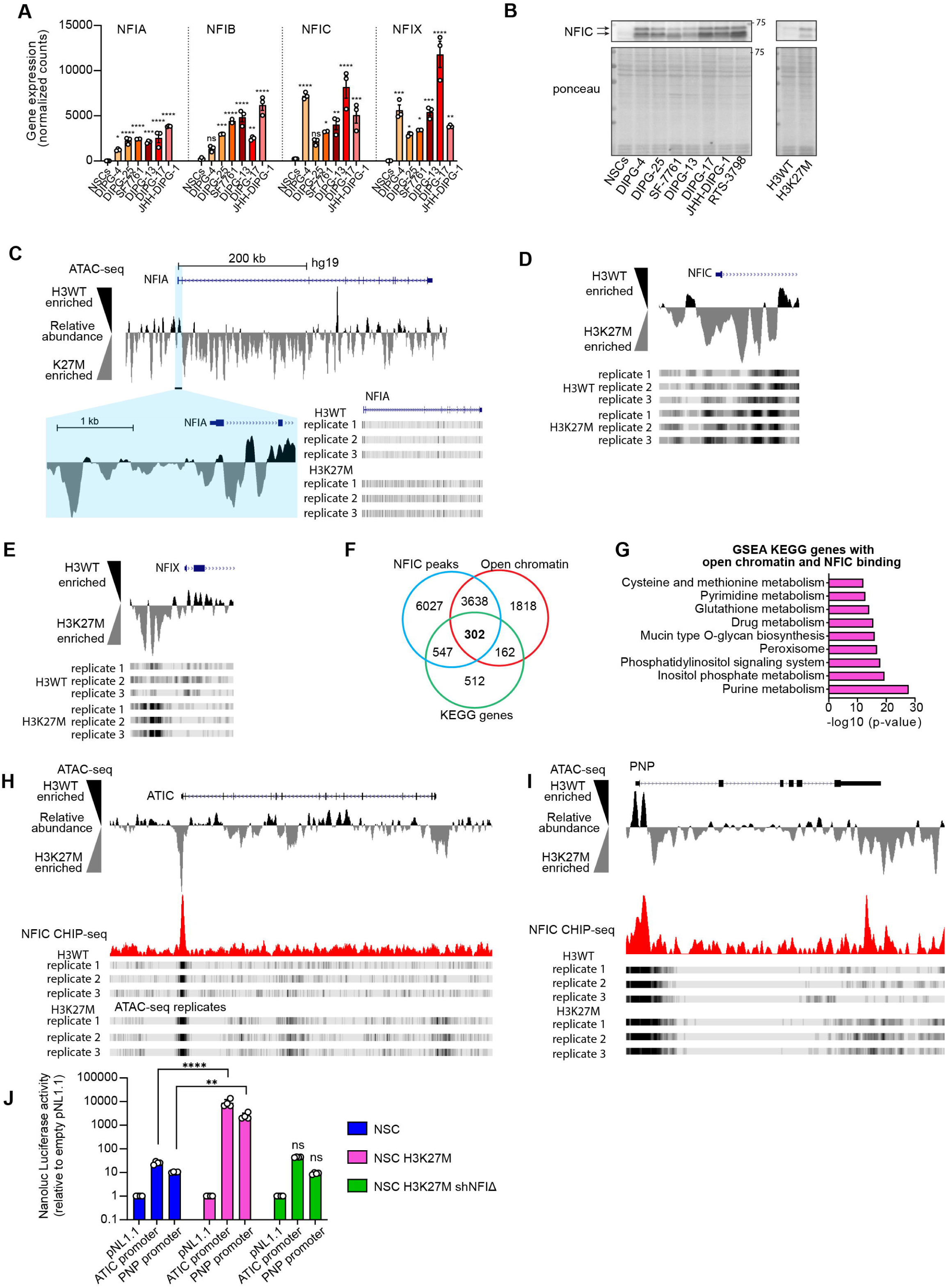
**NFI transcription factors mediate transcriptional activation of purine metabolism genes downstream of H3K27M mutation.** (A) Relative gene expression (normalized counts) of NFI-TFs from RNA-sequencing analysis (n=3). Each gene was analyzed separately by one-way ANOVA comparing DIPG lines to NSCs. (B) Western blot for NFI transcription factors in DIPGs vs NSCs and H3K27M NSCs vs H3WT NSCs. (C-E) ATAC-seq peaks at NFI loci showing increased chromatin accessibility in H3K27M NSCs. Data visualized using UCSC Genome browser with merge (subtract) overlay. H3K27M peaks are subtracted from H3WT peaks; positive (black) peaks indicate enrichment in H3WT, and negative (grey) peaks indicate enrichment in H3K27M. Individual replicates shown without merged overlay. (F) Venn diagram showing overlap of genes in KEGG pathways related to metabolism, and genes with more accessibility and strong NFIC binding in H3K27M-NSCs. Differentially accessible peaks (ATAC-seq) and NFIC peaks (ChIP-seq) in H3K27M-NSCs were assigned to genes using the Genomic Regions Enrichment of Annotations Tool (GREAT)^45,46^. (G) Enrichment analysis of KEGG genes that are more accessible in H3K27M-NSCs and have strong NFIC peaks (n=302 from panel M). (H) Merged ATAC-seq peaks at ATIC locus showing more accessibility at promoter/TSS (top) and strong NFIC binding determined by ChIP-seq (bottom) (I) ATAC-seq peaks at loci of PNP showing increased accessibility across gene body in H3K27M NSCs and strong NFIC binding at PNP promoter/TSS and intragenic enhancer region. (J) ATIC and PNP promoter activity measured by Luciferase reporter assay in NSCs, H3K27M-NSCs and H3K27M-NSCs with shRNA targeting NFI-TFs. For comparison between lines, the promoter activity was normalized relative to the mean of the luminescence units of the empty pNL1.1 vector. and C). Panel A plotted as mean +/- SEM, analyzed by one-way ANOVA (for each gene) with Dunnett’s correction. Panel J plotted as mean +/- SEM, analyzed by two-way ANOVA with Dunnett’s correction. (ns = not significant; * p < 0.05; ** p < 0.01; *** p < 0.001)

To determine whether NFI transcription factors are functionally linked to the mutation and altered gene expression in metabolic pathways, we performed ATAC-seq and ChIP-seq in isogenic normal human NSCs expressing wild-type or H3K27M histone 3 (n = 3 per group). Clear genotype-dependent clustering was observed across ATAC-seq replicates (Figure S4B). NFI genes demonstrated increased chromatin accessibility in H3K27M-mutant cells (Figures 4C–E,). Interestingly, motif enrichment analysis of regions with increased accessibility in H3K27M-NSCs revealed a strong overrepresentation of NFI binding motifs (Figure S4C).

An analysis of the peaks from NFIC ChIP-seq, performed in H3K27M NSCs, revealed gene targets related to developmental programs, chromatin regulation, and KEGG metabolic pathways (Supplemental Table S.7). NFIC target genes showed considerable overlap with regions of accessibility from ATAC-seq data (Figure 4F). Enrichment analysis with the overlapping genes that are in metabolic pathways (KEGG) were consistent with the transcriptomic and metabolomic alterations observed in DIPG (Figures 4G and S4D).

Notably, ATIC was not a predicted PRC2 target gene; however, NFIC ChIP-seq revealed strong binding at the ATIC promoter and showed increased chromatin accessibility by ATAC-seq for the H3K27M NSCs (Figure 4H). Similarly, the locus of the purine degradation gene PNP displayed broad accessibility across its gene body and prominent NFIC occupancy at the promoter region of the H3K27M expressing NSCs (Figure 4I).

Lastly, to test if the H3K27M mutation increases transcriptional activity of ATIC and PNP promoters in an NFI-dependent manner we cloned promoter regions of ATIC and PNP and performed luciferase activity assay in control or NFI-silenced H3K27M mutant NSCs. NSCs expressing H3K27M exhibited significantly higher luciferase activity driven by both promoters, which was suppressed upon shRNA-mediated silencing of all NFI TFs (Figure 4J and S4E-F). These results confirm that the H3K27M mutation alters chromatin structure, including increased accessibility to NFI-TFs and NFI-dependent regulation of metabolic pathways including purine metabolism, in DIPG.

### De novo purine biosynthesis is essential for DIPG growth and proliferation

Evidence of unregulated purine biosynthesis and degradation in DIPG suggested a potential vulnerability to inhibition of the de novo purine biosynthesis (DNPB) pathway. Gene silencing experiments were conducted using shRNAs targeting multiple DNPB enzymes (Figure S5A). All DIPG lines tested were sensitive to ATIC knockdown, even in the presence of excess hypoxanthine in the culture medium (Figures 5A–B, S5B), indicating that purine salvage alone is insufficient to sustain DIPG viability when DNPB is disrupted. Silencing of upstream enzymes, including GART and PAICS, also reduced cell viability in DIPG lines (Figures S5C–H). In H3K27M-expressing NSCs, which display elevated DNPB and degradation activity, ATIC knockdown led to a marked reduction in proliferation, whereas proliferation in wild-type H3 NSCs remained unaffected (Figure 5C, S5I), supporting that the H3K27M mutation sensitizes cells to DNPB loss.

**Figure 5.**
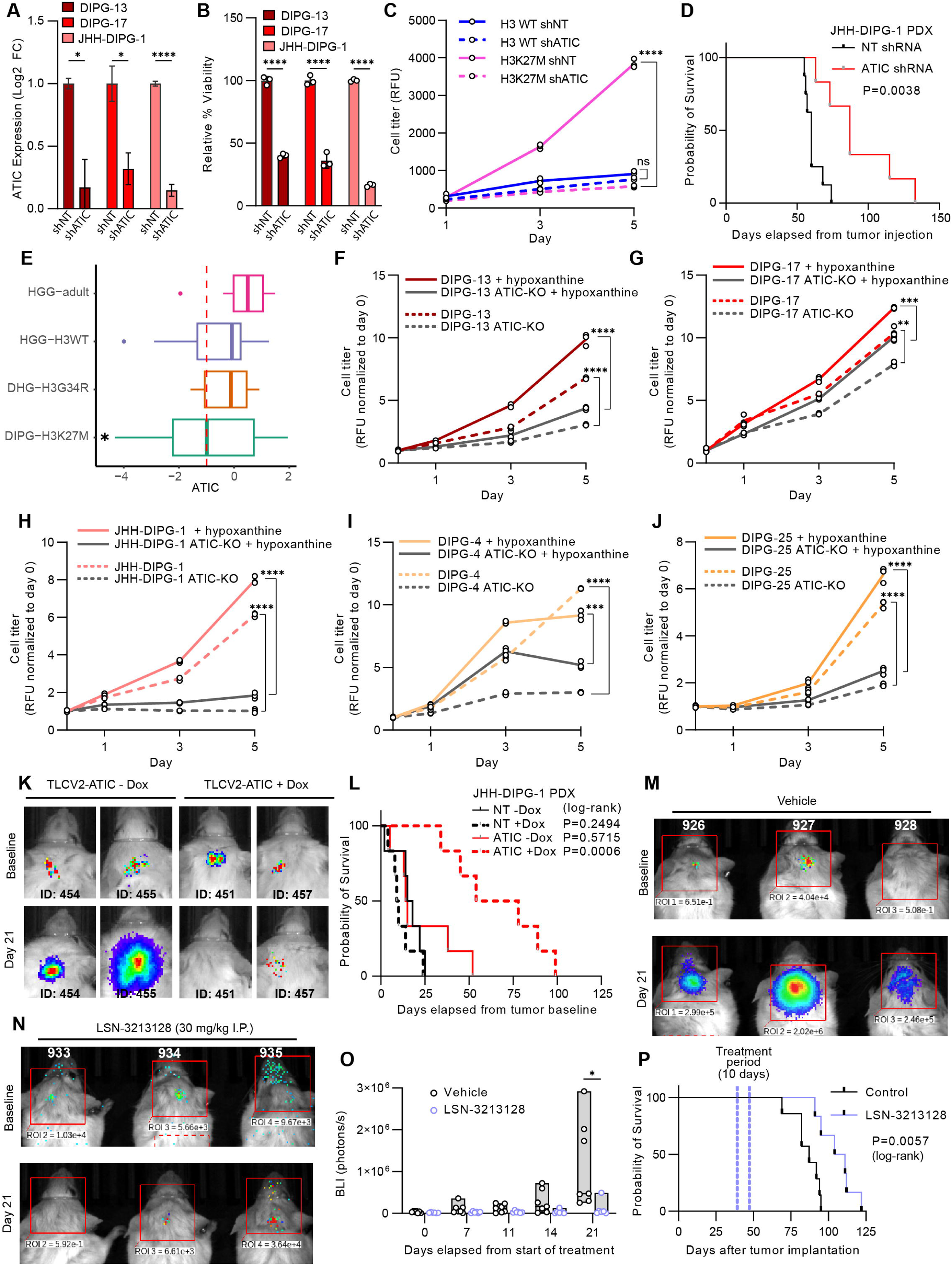
**De novo purine biosynthesis is essential for DIPG growth and proliferation.** (A) qRT-PCR showing fold change in mRNA levels in DIPGs stably expressing shRNA targeting ATIC relative to non-targeting (NT) controls (n=3). (B) Cell viability in DIPGs transduced with shRNA targeting ATIC relative to NT shRNA (n=3). Viability measured by CellTiter-Fluor Cell Viability Assay 72 hours after seeding in 96-well plate. (C) Proliferation of NSCs expressing H3WT or H3K27M, transduced with shRNA targeting ATIC or NT control (n=3), measured by CellTiter-Fluor (raw value plotted on y-axis). (D) Log-rank survival analysis for mice implanted with JHH-DIPG-1 cells stably expressing shRNA targeting ATIC (n=6) or NT control (n=8). (E) CRISPR dropout screen Box-whisker plot showing ATIC dependency in histone-mutation defined HGG (N=76) cell lines. HGG samples are grouped as HGG-adult (N=10, pink), H3 wildtype (WT) HGG (n=10, purple), H3K27M-DIPG (N=42 green) and H3G34 mutant DHGs (N=6, orange). The median is indicated by the thick line, the outer box shows the upper and lower quartiles. Whiskers show min/max values. Statistical significance was determined by Student t.test. * Denotes p value <=0.05. Red dash line at z score of −1 serves as a cut-off gene dependency. (F-J) Proliferation of DIPG lines stably expressing CRISPR gRNA targeting ATIC and a doxycycline-inducible CAS9(see also Figure S4J). Cells were plated 72 hours after induction of ATIC knockout in either normal growth media (15 µM hypoxanthine) or growth media supplemented with 35 µM hypoxanthine (50 µM total) (n=3). (K and L) Representative bioluminescent imaging (BLI) (K) and log-rank survival analysis (L) for mice implanted with luciferase-tagged JHH-DIPG-1 cells expressing doxycycline-inducible ATIC or NT CRISPR gRNA. Mice sorted into normal or doxycycline (dox) diet groups after confirming tumor by BLI (n=6) (see also Figure S4K for model and S4L for NT +/- dox). (M-P) Representative images (M and N), quantitation of BLI in region of interest (ROI) (O) and log-rank survival analysis (P) of mice implanted with JHH-DIPG-1 cells treated twice daily for 10 days with 30 mg/kg ATIC inhibitor LSN-3213128 (n=6) or vehicle control (n=7). BLI radiance expressed as total emission (photon/s) in a 2 cm x 2 cm red square (ROI shown in panel M and N). Panel A plotted as mean +/- SEM, and panel B plotted as mean +/- SD, analyzed by unpaired, two-tailed T tests. Panel C (day 5) analyzed by two-way ANOVA. Panels E-I (day 5) analyzed by multiple unpaired, two-tailed T tests. Panel M ROI quantitation plotted as floating bars with line at median, analyzed by two-way ANOVA with repeated measures.

Given the elevated oxidative stress observed in DIPG and the known role of purine metabolism in redox balance, the impact of ATIC depletion on reactive oxygen species (ROS) levels was evaluated. Although CRISPR-mediated ATIC knockout resulted in only a modest increase in mitochondrial ROS (12.3%) and no change in total ROS levels (Figures S5J–L), antioxidant supplementation with N-acetylcysteine (NAC) improved cell viability in both control and ATIC-depleted DIPG cells (Figures S5M–N). These results suggest that elevated baseline oxidative stress in DIPG may contribute to ATIC dependency; however, ATIC loss alone does not appear to further exacerbate ROS levels to a functionally significant extent.

To test the in vivo requirement for ATIC, two genetic systems were employed: ATIC shRNA and doxycycline-inducible CRISPR knockout. Following intracranial implantation of DIPG cells into postnatal day 2 (P2) NSG mice, ATIC shRNA expression significantly prolonged survival relative to nontarget controls (Figure 5D). For the dox-inducible CRISPR model, the essentiality for ATIC in DIPGs was first validated in vitro, and in parallel we investigated gene dependency data from the Childhood Cancer Model Atlas (CCMA). ATIC dependency is enriched in H3K27M-mutant DIPG relative to other high-grade glioma subtypes, including adult glioblastoma, H3G34R DIPG, and H3 wild-type DIPG (Figure 5E). CRISPR-Cas9 knockout of ATIC in five DIPG lines confirmed reduced viability across all models tested, which could not be rescued by hypoxanthine supplementation (Figures 5F–J, S5J), underscoring the essentiality of DNPB over salvage in DIPG.

Luciferase-labeled DIPG cells transduced with control or ATIC-targeting CRISPR were implanted intracranially into P2 NSG mice, and tumor burden was monitored following initiation of doxycycline diet to induce ATIC knockout. Tumor growth was substantially reduced in ATIC KO tumors (Figures 5K), compared to non-target control tumors (Figure S5O–P), and overall survival was significantly extended in the ATIC-targeted cohort (Figure 5L), confirming the therapeutic relevance of ATIC in vivo.

We next evaluated pharmacologic inhibition of ATIC. Compound 14 (Cpd14), a dimerization inhibitor that selectively targets the AICAR transformylase activity of ATIC, exhibited selective cytotoxicity in DIPG lines while sparing NSCs and NHAs (Figure S5Q). Despite its poor cell permeability and high micromolar effective dose, Cpd14 was well tolerated in mice at 50 mg/kg twice daily, and pharmacokinetic analysis confirmed brain penetration with measurable levels in plasma and brain extracellular fluid (Figures S5R–T). Treatment of DIPG-bearing mice with Cpd14 modestly extended survival (Figure S5U), demonstrating in vivo proof of concept.

To evaluate a potentially more translatable approach, the anti-folate LSN3213128 (LSN), which selectively targets the folate-dependent step of ATIC activity, was tested. A dosing schedule of twice daily IP injections at 30 mg/kg of LSN on a low-folate diet, based on previously published pharmacodynamic and anti-tumor studies[47], was well tolerated in preliminary toxicity studies (Figures S4V and W). LSN significantly attenuated tumor growth (Figures 5M–O) and extended overall survival (Figure 5P), despite the abbreviated dosing window. These results establish ATIC as a critical node in DIPG metabolism and highlight the therapeutic potential of targeting de novo purine biosynthesis in this disease.

### Adaptive resistance to ATIC inhibition recapitulates baseline dependency gene expression signatures related to dysregulated purine metabolism in DIPG

Although ATIC dependency is significantly enriched in H3K27M-mutant DIPG relative to other high-grade glioma subtypes (see Figure 5E), the degree of dependency varies considerably among H3K27M lines, with a wide range of dependency z-scores. Given this variability, we investigated whether gene expression signatures are associated with differential ATIC dependency to identify features that provide further insight into the molecular mechanisms driving ATIC sensitivity and resistance.

A stable ATIC inhibitor–resistant DIPG model was generated by continuously passaging one of the most ATIC-dependent cell lines, JHH-DIPG-1 (ATIC z-score = −3.024), in increasing concentrations of the ATIC inhibitor Cpd-14 (Figure 6A). Cell viability assays confirmed a marked reduction in Cpd-14 sensitivity in the resistant cells compared to the parental line (Figure S6A), validating successful acquisition of resistance.

**Figure 6.**
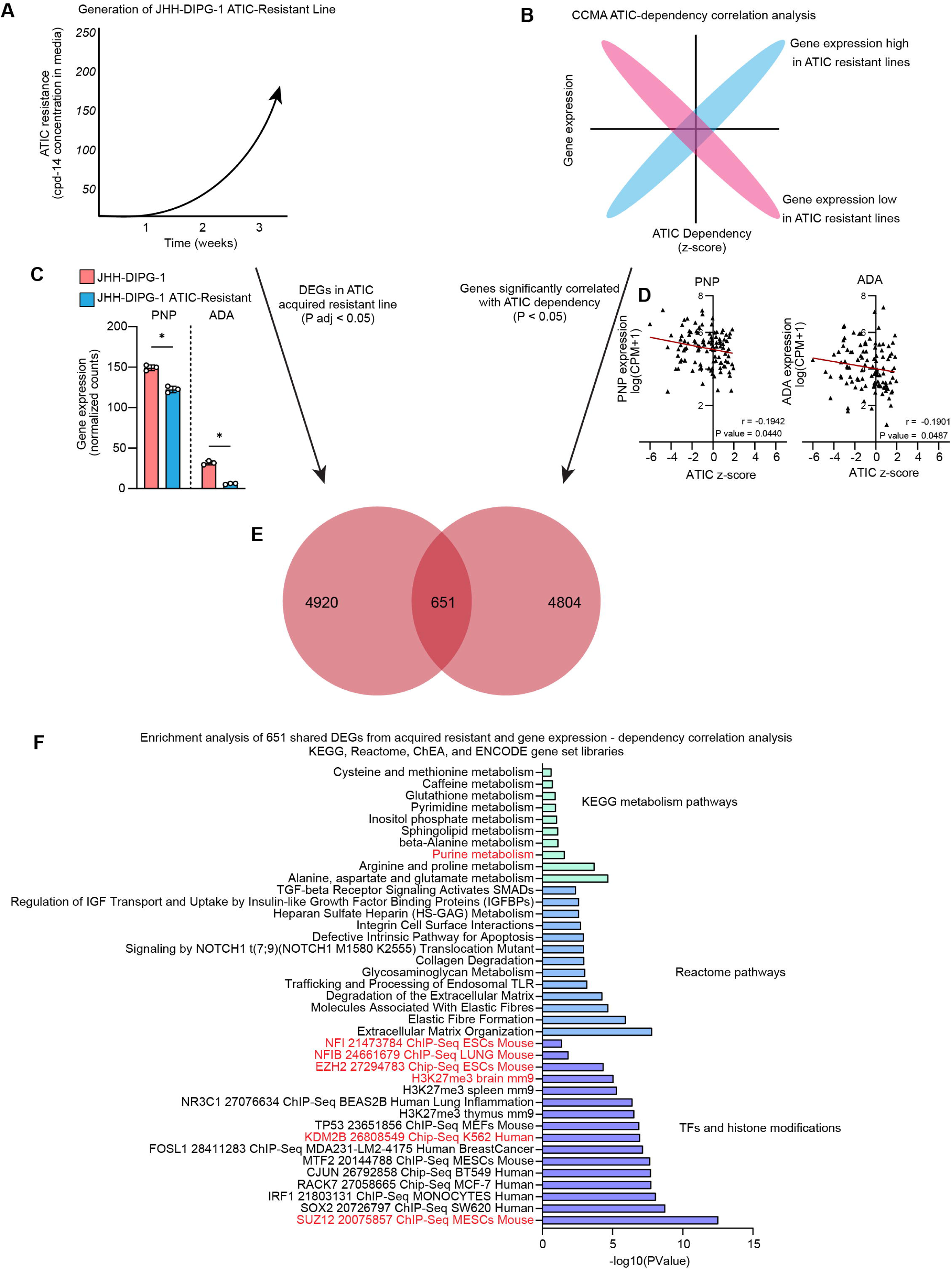
**Adaptive resistance to ATIC inhibition recapitulates baseline dependency gene expression signatures related to dysregulated purine metabolism in DIPG** (A) Schematic overview for the generation of an ATIC inhibitor–resistant DIPG line established by prolonged exposure to Compound 14 (Cpd14). (B) Schematic for ATIC-dependency vs gene expression correlation analysis to identify genes upregulated or downregulated in ATIC-knockout resistant cell lines from the Childhood Cancer Model Atlas (CCMA) CRISPR screening dataset. (C) Bar plot of gene expression (normalized counts) for purine degradation enzymes PNP and ADA in JHH-DIPG-1 parental and ATIC acquired resistant line. (D) Scatter plot of ATIC z-score and gene expression of PNP and ADA in CCMA cell lines (E) Venn diagram comparing differentially expressed genes in the acquired resistant model (Padj < 0.05, n = 5,571) and genes that significantly correlated with ATIC dependency in the CCMA dataset (p < 0.05, n = 5,455). Overlap of 651 genes was identified with concordant directionality. (F) Enrichment analysis of the 651 overlapping genes (From 6E), using KEGG, Reactome, ChEA, and ENCODE databases. Highlighted enriched gene sets include PRC2 targets (*EZH2*, *SUZ12*), histone modification H3K27me3, purine metabolism, and transcriptional regulators such as nuclear factor I (NFI) family.

To identify molecular features associated with ATIC resistance, we compared gene expression changes in the resistant line with baseline gene expression features linked to ATIC dependency across all models in the Childhood Cancer Model Atlas (CCMA). Specifically, we performed RNA-seq on parental and resistant JHH-DIPG-1 cells and identified 5,571 differentially expressed genes (DEGs) (adjusted p < 0.05). Separately, a correlation analysis between gene expression and ATIC dependency z-scores was conducted using CRISPR screening data from the CCMA dataset (Figure 6B), revealing 5,455 genes whose expression significantly correlated with ATIC dependency (p<0.05).

Comparison of these two datasets uncovered 651 genes significantly altered in both contexts, including concordant downregulation of purine metabolism enzymes involved in DNPB and degradation (Figure 6C-E, S6B and C; see also Supplemental Table S.9). Notably, PNP and ADA, key enzymes in purine catabolism, and the DNPB genes PFAS and ADSL were downregulated in the ATIC-resistant line and showed negative correlation with ATIC dependency in the CCMA correlation analysis.

We performed enrichment analyses on the 651 genes that were both differentially expressed in the ATIC-resistant DIPG line and significantly correlated with ATIC dependency in the CCMA dataset (Figure 6F). Gene set enrichment revealed strong overrepresentation of transcriptional and chromatin-based regulatory programs related to the pathobiology of DIPG, including EZH2 and SUZ12 target genes, as well as genes marked by H3K27me3, implicating re-engagement of PRC2-mediated repression in the ATIC-resistant state. Consistent with upstream findings, targets of the NFI-TFs were also significantly enriched, reinforcing their role related to altered metabolism and epigenetic reprogramming in DIPG.

These findings demonstrate that suppression of unregulated purine biosynthesis and degradation is part of the adaptive transcriptional program associated with resistance to ATIC inhibition, potentially serving to conserve purine pools or prevent toxic accumulation of degradation byproducts under metabolic stress. This reinforces our earlier results showing that dysregulated purine metabolism is a consequence of H3K27M mutations, driven in part by aberrant NFI transcriptional activity. Together, these data establish de novo purine biosynthesis as a therapeutic vulnerability in DIPG and highlight the convergence of metabolic adaptation and epigenetic repression as defining features of acquired resistance to ATIC inhibition.

## DISCUSSION

Diffuse intrinsic pontine glioma (DIPG) remains one of the deadliest pediatric cancers, with limited treatment options and dismal survival outcomes. At the center of its pathogenesis is the H3K27M mutation, which drives epigenetic dysregulation through PRC2 inhibition and altered histone methylation^28,29^. While the mutation’s impact on gene expression is well characterized^30,31^ its consequences on metabolism remain less defined. This study identifies dysregulated purine metabolism as a defining metabolic feature of H3K27M-mutant DIPG. Using a multi-omics approach across human, murine, and xenograft models, we show that DIPGs exhibit simultaneous upregulation of de novo purine biosynthesis (DNPB) and purine degradation, resulting in elevated levels of inosine, hypoxanthine, and downstream catabolic products. This phenotype is driven by aberrant upregulation of the rate-limiting enzyme ATIC and reflects a unique form of metabolic reprogramming downstream of H3K27M mutations.

The simultaneous upregulation of purine biosynthesis and unregulated purine degradation, resulting in elevated purine pools and their waste products are in line with previous reports showing elevated purine pools in DIPGs and normal cells expressing H3K27M^22,32^. Notably, our results are also consistent with a recent study where enrichment of purine nucleotides was observed in a few DIPG lines that were maintained in a stem cell like state^33^. While no mechanisms for purine enrichment were examined in this report, we are the first group to demonstrate how a DIPG-specific mutation (H3K27M) causally regulates purine metabolism (and other pathways) through transcription factors (NFI) that are yet unknown to regulate purine metabolism in brain tumors. Furthermore, the recent report on DIPG addiction to de novo pyrimidine biosynthesis is mechanistically similar, where the authors show the dependence on pyrimidine biosynthesis is directly related to an upregulation of pyrimidine degradation^27^. Our findings also reveal unregulated purine metabolism in DIPG is linked to elevated oxidative stress, suggesting a possible feedforward loop in which purine degradation contributes to oxidative stress, which in turn may reinforce continued purine catabolism.

Through our investigation into the regulatory mechanisms, we identified a relationship between NFI-TFs and the expression of genes related to the altered metabolic phenotype. H3K27M mutation was sufficient to induce purine metabolic rewiring through NFI family transcription factors, which were consistently upregulated and identified as direct regulators of ATIC and purine degradation genes. Integration of ATAC-seq and NFIC ChIP-seq data confirmed increased chromatin accessibility and transcriptional control at key purine gene loci, reinforcing the link between epigenetic reprogramming and altered metabolism in DIPG. While other pediatric cancers show dependency on nucleotide metabolism, these findings establish a direct molecular mechanism linking an oncogenic histone mutation to purine pathway activation via a defined transcriptional axis. NFI-TFs were identified as PRC2 targets in multiple curated datasets. Further, NFIC has been associated with super-enhancers in DIPG, and its expression is reduced following treatment with a histone deacetylase (HDAC) inhibitor^31^. While NFI-TFs were not thematically explored in prior studies, data from isogenic CRISPR-edited H3K27M-WT DIPG cell lines—including differential expression, ATAC-seq, and regulatory network analyses— indicate that NFI family members are upregulated in the H3K27M context and are enriched at accessible chromatin regions linked to neurogenesis, cell fate, and stemness pathways as super-enhancers^34,35^. Although these findings were not the focal point of the original papers, they support a model in which NFI transcription factors contribute to the transcriptional reprogramming downstream of H3K27M-mediated epigenetic dysregulation. This underscores the pivotal role of epigenetic changes driven by H3K27M mutations influencing transcriptional network changes, with NFI-TFs playing a central role. The high expression of these genes in DIPG cell lines was concordant with what we saw in neural stem cells expressing the H3K27M mutation and validated by our CHIP-sequencing data. Although other mechanisms for purine enrichment were not directly addressed in this study, we provide the first evidence that the H3K27M mutation causally reprograms purine metabolism in DIPG through NFI transcription factors—regulators not previously linked to purine control in brain tumors. This expands upon prior work implicating MYC in driving de novo purine synthesis in brain tumor-initiating cells (BTICs)^36^, and complements previous findings that MYC activation downstream of H3K27M also contributes to broader metabolic rewiring in DIPG^37^. Together, these studies suggest that metabolic dependencies in DIPG can arise through distinct, mutation-driven transcriptional programs, with NFI representing an alternative axis to MYC. These transcriptional network changes are characteristic of DIPGs in the context of their precursor cells and should be further investigated.

We investigated the therapeutic potential of targeting DNPB and demonstrated that DIPG cells are highly dependent on this pathway. Knockdown or knockout of ATIC significantly impaired viability across DIPG lines and extended survival in orthotopic mouse models. This dependency could not be rescued by hypoxanthine supplementation, indicating limited capacity for salvage under nutrient-limited conditions of the brain microenvironment, where purine levels are known to be lower^38–41^. Notably, a recent study showed that H3K27M-mutant DMGs can upregulate purine salvage in response to radiation, particularly via guanine salvage pathways^42^, suggesting that context-dependent metabolic plasticity may shape resistance mechanisms depending on the stressor. In contrast, our findings reveal that salvage is insufficient to rescue ATIC loss, highlighting a critical vulnerability in DNPB that may be therapeutically exploitable in settings where salvage is constrained.

Pharmacological inhibition of ATIC with two different compounds reduced tumor proliferation and increased overall survival in a mouse model of DIPG. LSN-3213128, a non-classical antifolate that selectively targets ATIC^43,44^, had a robust anti-tumor effect. The dosing regimen was paired with a folate-free diet, consistent with prior reports showing enhanced activity of LSN under low-folate conditions^43^. Although these results were encouraging, a key limitation of our study was the abbreviated duration of drug administration, necessitated by limited compound availability and high cost, which may have constrained the full therapeutic potential of LSN in vivo.

Although LSN was more effective in vivo, we investigated mechanisms of resistance using the ATIC homodimerization inhibitor Cpd-14, which does not rely on folate transport or retention. This choice was critical, as the antifolate structure of LSN could potentially engage canonical antifolate resistance mechanisms—such as altered transporter expression or folate pool competition—that might obscure resistance mechanisms specific to ATIC inhibition.

Long-term exposure to ATIC inhibitors resulted in suppression of purine metabolic gene expression and reactivation of PRC2-mediated repression. Downregulation of both DNPB and purine degradation genes in resistant cells closely mirrored gene expression signatures observed in tumors with low ATIC dependency, indicating a metabolic shift toward repression as an adaptive mechanism of resistance.

These findings raise the possibility that PRC2-mediated repression in DIPG may be metabolically gated, responding to the bioavailability of key metabolites consumed by DNPB such as glycine, that we show to be low DIPGs (from Supplementary Table S.1). This is consistent with prior work showing that α-KG availability, shaped by glucose and glutamine metabolism, regulates H3K27me3 levels in H3K27M gliomas^22^ and suggests that purine metabolic rewiring may participate in feedback loops influencing chromatin state. Overall, our research highlights the significant links between epigenetic mutations and metabolic changes in DIPG, providing valuable insights for future treatment approaches for this lethal childhood cancer.

## Data storage

The information on the sequenced and partially processed RNASeq datasets, ATACseq datasets and ChIP-Seq datasets have been deposited to the National Center for Biotechnology Information (NCBI)’s GEO database. The GEO accession number to access the datasets will be provided.

## ACKNOWLEDGEMENTS

Our sincere thanks to Michelle Monje who provided the majority of the DIPG cell lines, and to Eric Raabe and Rintaro Hashizume for providing the remaining DIPG cell lines. We would like to thank Cincinnati Children’s core facilities including genomics, flow cytometry and viral vector core facilities, and the University of Colorado SOM Metabolomics core facility. The University of Colorado SOM Metabolomics core acknowledges support from the University of Colorado Cancer Center (P30CA0469340). This work was supported by NIH P30 AR070549 and Cincinnati Children’s Hospital ARC Award #53632 to M.T.W. and L.K.; a Cancer Research UK grant to A. T. (A20185); Pilot Innovation Award from Cincinnati Children’s, multiple awards from CancerFreeKids, CureStartsNow and TeamConor Childhood Cancer Foundation, and awards from National Institute of Health (R01 NS132884 (to B.D.)

## AUTHOR CONTRIBUTIONS

I.M. and B.D. conceived the experiments, analyzed data, and wrote the manuscript. I.M. performed most of the experiments and prepared figures. SC., M.H. R.C., R.M and J.D performed additional experiments. S.C, M.H. and R.C assisted in in vivo and in vitro experiments. J.D, and L.S. performed and P.D. designed PK studies, and J.A.R and A.G. performed and A.D. supervised metabolomics studies. M.R. H and S.C designed and performed luciferase promoter activity assay. O.D. performed ATAC-seq, and ChIP-Seq experiments and M. T. W. and L.K. designed ATAC-seq, and ChIP-Seq experiments, analyzed data and assisted in manuscript writing. C.S performed whole genome CRISPR studies in cancer cell lines and R.F supervised this study and assisted in manuscript writing. C.B.S, P.D, N.P.S, and T.H provided clinical samples, N.E. and A.T. provided Compound 14 and T.P provided DIPG GEMM.

## DECLARATION OF INTERESTS

AT is a director and shareholder of Curve Therapeutics, who are developing Compound 14 towards the clinic.

## INCLUSION AND DIVERSITY

We support inclusive, diverse, and equitable conduct of research.

## RESOURCE AVAILABILITY

### Lead contact

Further information and requests for resources and reagents should be directed to and will be fulfilled by the lead contact, Biplab Dasgupta.

### Materials availability

All unique reagents generated in this study are available from the lead contact without restriction.

### Data and code availability

RNA-seq, ATAC-seq, ChIP-seq and Metabolomics data are being deposited in GEO and publicly available as of the date of publication. Accession numbers will be listed in the key resources table. Any additional information required to reanalyze the data reported in this paper is available from the lead contact upon request.

## SUPPLEMENTAL FIGURE LEGEND

**Figure S1. Purine metabolism is consistently dysregulated across human, murine, and PDX models of DIPG, related to Figure 1**.

(A and B) GSEA from RNA-seq data (A) and Metaboanalyst joint-pathway analysis integrating RNA-seq and metabolomics data (B) comparing two distinct groups of DIPG cell lines.

A. Enrichr gene set enrichment plots for gene sets in ARCHS4_Kinases_Coexp and ENCODE_and_ChEA_Consensus_TFs_from_ChIP-X databases from DEGs in 6 DIPG cell lines vs human NSCs (n=3 each).

(D-E) Metaboanalyst enrichment analysis and joint-pathway analysis for differentially expressed genes and differentially regulated metabolites in DIPG IUE mouse model cell lines relative to normal mouse brainstem NSCs.

(F) Relative abundance of purine degradation products in DIPG IUE model relative to control mouse brainstem NSCs (n=9 each, 3 separate lines derived from tumors or normal brainstem performed in triplicate).

(G) PLS-DA of metabolites in DIPG-PDX vs normal brainstem.

(H) Relative abundance (sum normalized peak area) of purine nucleotides and nucleosides from DIPG tumors in mice. Tumors derived from two separate DIPG cell lines individually compared to normal mouse brains (n=5).

(I) Gene expression of purine cyclase and phosphodiesterase enzymes involved in purine nucleotide conversion to second messengers.

(J-R) Relative abundance (sum normalized peak area) of metabolites in cell lines (n=6) (J-L), DIPG PDX tumors (n=5) (M-O), and mouse IUE lines (n=9) (P-R). F,H and J-R plotted as mean +/- SEM analyzed by unpaired, two-tailed T test (ns = not significant; * p < 0.05; ** p < 0.01; *** p < 0.001; **** p < 0.0001).

**Figure S2. Dysregulated purine metabolism and high oxidative stress in DIPG, related to Figure 2**.

(A , B) Tracing analysis of isotope-labeled serine and that of hypoxanthine (C, D) after 6-h incubation showing fraction of labeled metabolites detected in purine nucleotides (n=3).

(E) Relative abundance (peak area) of labeled inosine from 13C6-glucose tracing analysis. (F-H) 5-hydroxyisourate detected at higher levels in DIPGs across experiments (n=3).

(I-L) Relative gene expression (normalized counts) for genes related to purine salvage and degradation from RNA-sequencing analysis (n=3). Each gene was analyzed separately by one-way ANOVA comparing DIPG lines to NSCs.

(M) Ratio of peak areas for reduced glutathione (GSH) and oxidized glutathione (GSSH) as a marker for oxidative stress, from untargeted metabolomics in Figure 1A (n=6).

(N-P) Dehydroascorbate (unlabeled) detected across multiple experiments at higher levels in DIPGs as another marker of oxidative stress (n=3).

(Q) Gene expression of ROS related genes in pediatric low-grade glioma (LGG), high-grade glioma with wildtype Histone 3 (HGG H3WT), and diffuse midline glioma with Histone 3 K27M mutation (DMG H3K27M). Data obtained from PedcBioPortal’s OpenPBTA dataset^47^.

(R) Relative gene expression (log2 fold change) for ROS related genes in DIPG cell lines relative to NSCs.

(S and T) Flow cytometry analysis of total (CellROX Orange) and mitochondrial (MitoSOX Red) ROS levels in DIPGs and NSCs.

Panels A - D plotted as mean +/- SEM, analyzed by one-way ANOVA with multiple comparisons. PanelsE-H and N-P plotted as mean +/- SEM. Each DIPG line was compared to NSCs, analyzed by unpaired, two-tailed T test. Panels I-L plotted as mean +/- SEM, analyzed by one-way ANOVA (for each gene) with Dunnett’s correction. Panel M plotted as mean +/- SEM analyzed by one-way ANOVA. (ns = not significant; * p < 0.05; ** p < 0.01; *** p < 0.001; **** p < 0.0001)

**Figure S3. H3K27M mutation drives dysregulated purine metabolism through upregulation of ATIC and purine pathway genes, related to Figure 3**.

(A-F) Gene expression of DNPB genes and degradation enzyme PNP (FPKMs) in H3WT patients (n=1786) and H3K27M patients (n=176) from pediatric brain tumor atlas (PedcBioportal).

(G) Log-rank and Gehan-Breslow-Wilcoxon survival analysis in DIPG patients stratified by high ATIC expression (above median, n=16) and low ATIC expression (below median, n=16).

(H) Log-rank survival analysis in pediatric brain tumor atlas (PedcBioportal, PBTA); patients stratified by high ATIC expression (above median, n=428) and low ATIC expression (below median, n=428).

(I) Representative IHC showing ATIC protein expression in DIPG-PDX mouse model (n=3). Tumor tissue and adjacent normal tissue marked by boundary.

(J-K) Normalized counts for DNPB (J) and purine degradation genes (K) from RNA-seq data in tumors from DIPG IUE mouse model. This model used PdgfraD842V, dominant negative Trp53 (DNp53), and H3.3K27M which induced fully penetrant brainstem gliomas with histopathological and molecular features found in human DIPGs. RNA-seq data downloaded from Gene Expression Omnibus GSE128807.

(L) Western blot comparing protein expression levels of ATIC in normal mouse NSCs relative to IUE-24B1 (cell line derived from DIPG GEMM^48^).

Panels A-F plotted as scatter dot plot with line at median, analyzed by two-tailed T test with Welch’s correction. Panels J and K plotted as mean +/- SEM, analyzed by DESeq2 pipeline.

**Figure S4. NFI transcription factors mediate transcriptional activation of purine metabolism genes downstream of H3K27M mutation, related to Figure 4**.

(A) qRT-PCR showing fold change in NFI-TFs mRNA levels in NSCs expressing H3K27M relative to NSCs expressing H3WT (n=3).

(B) Clustering analysis of ATAC-sequencing replicates shows strong agreement between replicates.

(C) Regions of chromatin accessibility were examined for transcription factor binding site motifs, showing NFI-TF motif enrichment in regions with more accessibility identified in H3K27M NSCs.

Overlap of differentially expressed KEGG genes in DIPGs (relative to NSCs) and KEGG genes in H3K27M-NSCs, with open chromatin and strong NFIC binding (NFIC target genes).
Western blot for antibodies specific to NFIC (left) or all NFI-TF isoforms (right) in H3K27M-NSCs transduced with shRNAs targeting all NFI-TFs or shRNAs targeting only NFIC.

Panel A plotted as mean +/- SEM, each gene compared to H3WT NSCs analyzed by unpaired, two-tailed T test.

**Figure S5. De novo purine biosynthesis is essential for DIPG growth and proliferation, related to Figure 4**.

(A) Western blot showing shRNAs screened for knockdown efficiency of DNPB genes.

(B-D) Proliferation of JHH-DIPG-1 cells transduced with shRNA targeting DNPB genes ATIC (B), PAICS (C), and GART (D) or NT control (n=3), measured by CellTiter-Fluor (raw value plotted on y-axis).

(E-H) Relative expression measured by qRT-PCR for DNPB genes PAICS (E) and GART (G) in two additional DIPG lines (DIPG-13 and DIPG-17) transduced with shRNA targeting these genes relative to NT control, and cell viability in DIPGs expressing shRNA targeting PAICS (F) and GART (H) relative to NT control (n=3).

(I) Western blot showing ATIC knockdown in H3WT and H3K27M NSCs transduced with shRNA targeting ATIC or NT control.

(J) Representative western blot for TLCV2 mediated doxycycline-inducible CRISPR knockout of ATIC.

(K and L) Representative flow cytometry analysis of mitochondrial (MitoSOX Red) and total (CellROX Orange) ROS levels in DIPGs with or without ATIC-KO.

(M and N) Flow cytometry analysis of ROS levels (M) and cell viability in DIPGs, with or without ATIC-KO, treated with 1 mM NAC.

(O) Schematic of in vivo TLCV2-ATIC knockout in luciferase-tagged JHH-DIPG-1 cells implanted into the pons of mice. IVIS imaging was performed twice weekly until tumors were detected, then mice were sorted into +/- dox groups.

(P) Representative images of mice implanted with JHH-DIPG-1 cells expressing TLCV2 with NT gRNAs on dox diet showing no inhibition of tumor growth relative to normal diet.

(Q) Dose response curve for NSCs, NHAs, and DIPGs, treated with increasing concentrations of ATIC dimerization inhibitor cpd-14.

(R) PK studies on cpd-14 show concentration of cpd14 at 50 mg/kg i.p. in NSG mouse brain (n=2/time point) at indicated time points followed by M/S detection.

(S and T) Plasma (S) and ECF (T) cpd-14 concentrations in rats (n=6) measured by LC-MS.

(U) Log-rank survival analysis of mice implanted with JHH-DIPG-1 cells treated with 50 mg/kg twice daily of cpd-14 (n=14) or vehicle control (n=20).

(V and W) Toxicity studies on mice treated with 30 mg/kg twice daily LSN-3213128 for 10 days or vehicle control on a low-folate diet shown by complete blood count (CBC) analysis (n=3) (V) and body weight throughout treatment (n=6) (W).

Panels B-D and Q plotted as mean +/- SD. Panels E-H and N plotted as mean +/- SD analyzed by unpaired, two-tailed T tests.

**Figure S6. Adaptive resistance to ATIC inhibition recapitulates baseline dependency gene expression signatures related to dysregulated purine metabolism in DIPG, related to Figure 6**

(A) Cell viability in JHH-DIPG-1 parental and ATIC acquired resistant line treated with increasing concentrations of cpd-14.

(B and C) Scatter plot of ATIC z-score and gene expression for DNPB genes PFAS and ADSL in CCMA cell lines.

## EXPERIMENTAL MODEL, MATERIALS AND METHODS

### Cell Culture

DIPG lines were obtained from our collaborators; (SU-DIPG-IV, SU-DIPG-XIII, SU-DIPG-XVII, SU-DIPG-XXV were provided by Dr. Michelle Monje, JHH-DIPG1 cells were provided by Dr. Eric Raabe at Johns Hopkins University; SF7761 cells were provided by Dr. Rinataro Hashizume and Dr. C. David James at Northwestern University and all have previously been described^49–51^ RTS-3798 is a newly-established DIPG line in our lab, derived by surgical biopsy from a DIPG patient in our laboratory. Mouse NSCs and IUE lines, derived from IUE mouse model of DIPG previously described^48^, were provided by Dr. Timothy Phoenix. All human cell cultures were generated with informed consent and in compliance with Institutional Review Board (IRB)-approved protocols. Cells were maintained as suspension cultures in UltraLow attachment plates in glioma stem cell (GSC) medium that contained DMEM-F/12 supplemented with B27, EGF (10 ng/ml), bFGF (10 ng/mL) and heparin (5 mg/mL).

Normal human astrocytes (NHA) were purchased from Lonza Group Ltd. and immortalized with retroviral expression of Large T-antigen (pBabe-puro TcDNA, #14088, Addgene). NHA was maintained in DMEM-F/12 supplemented with 10% FBS.

Human Neural Stem Cells (H9 hESC-Derived) were purchased from Gibco and maintained in GSC media, grown as adherent monolayer on Geltrex coated plates.

All lines were routinely checked for mycoplasma every other week using mycoplasma-specific PCR.

### Untargeted metabolomics in 6 DIPG cell lines and NSCs

DIPG cell lines and NSCs were plated in 10 cm dishes (n=6 each). ∼1 x 10^6^ cells were collected, washed with ice-cold PBS, centrifuged and flash frozen using liquid nitrogen. Untargeted metabolomic profiling was performed at Mayo using an Ultra HPLC (Infinity 1290 UHPLC, Agilent Inc., U.S.A.) coupled to 6550 quadruple ToF MS (Q-ToF MS, Agilent Inc., U.S.A.). Data were acquired both in positive and negative ESI modes in the mass range of m/z 100–1600 at a resolution of 10000. Chromatographic separation was achieved using hydrophilic interaction LC (HILIC) and reverse phase (C18) LC separately.

Metabolomics data processing and analysis (from untargeted metabolomics in cell lines)

All raw data files were converted into compound exchange file (CEF) format using masshunter profinder (version B08.00) software (Agilent). Mass profiler professional (Agilent Inc., U.S.A.) was used to convert metabolite features from each data file into a matrix of detected peaks for compound annotations and statistical analysis. ID Browser within the mass profiler professional (MPP) software v13.1 (Agilent) was used to annotate metabolites using data from METabolite LINk (METLIN), human metabolome database (HMDB), Kyoto Encyclopedia of Genes and Genomes (KEGG), and lipid maps. Annotations are based on isotope ratios, accurate mass, chemical formulas, and database score. Statistical analysis and enrichment analysis was performed in MetaboAnalyst and assigned to KEGG curated pathways.

### CRISPR screen analysis

Gene dependency scores for ATIC1 were analyzed from pooled CRISPR/Cas9 knockout screens conducted by the (CCMA) program from a panel of 76 high grade glioma cell lines using a targeted library of 352 cancer related genes and controls^52^. Briefly, Cas9-engineered cell line models were transduced with a pooled sgRNA library and grown for 21-day to assess effects on cell growth. Gene-level Z score was determined using the maximum likelihood estimation algorithm using the MAGECK package^53^.

### RNA-Sequencing in DIPGs and NSCs

∼ 1 microgram of RNA was prepared for each line (RNeasy kit , Qiagen) and used for mRNA library preparation. Library preparation and sequencing on the Illumina NovaSeq6000 were performed by the CCHMC DNA Sequencing and Genotyping Core.

Differential gene expression analysis performed using Deseq2 pipeline^54^. Briefly, HISAT2 was used on fastq files for alignment to reference genome HG38^55^ and Featurecounts was used on BAM files to produce counts data^56^. DESeq2 was used to determine differentially expressed genes and normalized counts tables for comparison between all 6 DIPGs to NSCs. Differentially expressed genes were analyzed using GSEA with KEGG gene sets^57,58^.

### DIPG mouse model and in vivo metabolomics

All animal procedures were carried out in accordance with the Institutional Animal Care and Use Committee (IACUC)-approved protocol of Cincinnati Children’s Hospital Medical Center (CCHMC; Cincinnati, OH). Animals were monitored daily by veterinary services.

For in vivo metabolomics analysis 100,000 JHH-DIPG-1 or SU-DIPG-XVII cells were injected into cold-anesthetized, postnatal day 2 mouse pups by stereotactic injection through a 31G burr hole (stereotactic coordinates: 3 mm posterior to lambda suture and 3 mm deep).

For in vivo bioluminescent imaging, luciferase expressing cells were established by infection with plenti-CMV-luc viral particles (a gift from Dr. Susanne Wells) and tumor growth was monitored using IVIS Spectrum CT system. Five minutes before bioluminescence imaging, mice were anesthetized and injected (intraperitoneally) with luciferin (150 mg/kg). All mice were euthanized following observation of lethargy and/or neurologic symptoms.

After confirming the approximate location of brain tumors via IVIS imaging, mice were sacrificed by isoflurane overdose. After rapid decapitation, brains were immediately harvested and brainstem tissue of tumor and non-tumor bearing mice was flash-frozed with liquid-nitrogen and stored at −80^0^C.

Metabolomics analyses were performed at the University of Colorado School of Medicine Metabolomics Core (Aurora, CO). Metabolites were extracted from frozen tissue as previously described^59^ for analysis by UHPLC-MS (Thermo Vanquish UHPLC/Thermo Q Exactive mass spectrometer (Thermo Fisher Scientific, Waltham, MA, USA)). Samples were run both in positive and negative ion mode (separate runs) using 1 min or 5 min gradient methods as previously described^59^. To account for uncertainty in cell number per pellet, total protein quantity was used to normalize metabolite abundances. For protein quantification, each post-extraction cell pellet was resuspended in PBS at the same approximate concentration (2million cells/mL) as during metabolite extraction. Each sample was then measured for UV absorbance (λ = 280 nm, Nanodrop, Thermo Scientific, Waltham, MA USA) in triplicate and metabolite abundances were normalized to the average of these three replicates.

Mass spectral features were identified, quantified by peak area and annotated using El-Maven^60^ v0.12.0 with the KEGG database. MetaboAnalyst 5.0 was used for statistical analysis (PLS-DA, Heatmaps) and pair-wise comparison of sample groups^61^.

### Stable-isotope tracing with 13C6-glucose, 15N-serine, and 15N-hypoxanthine

DIPG cell lines and NSCs were incubated for 6 hours in media containing stable-isotopes before being washed with ice-cold PBS, centrifuged and flash-frozen. For 13C6-glucose tracing, glucose-free DMEM was supplemented with [^13^C_6_]glucose (Cambridge Isotope) for a final volume of 5mM. For ^15^N-Serine tracing and ^15^N_4_-hypoxanthine tracing, MEM was used supplemented with either 60 μM unlabeled hypoxanthine and 60 μM 15N Ser, or 60 μM ^15^N_4_ Hypo and 60 μM unlabeled Ser respectively, as previously described^62^. Metabolite extraction and profiling performed on UHPLC[MS using a Vanquish UHPLC coupled to a Q Exactive mass spectrometer as previously described. Mass spectral features were identified, quantified by peak area and annotated using El-Maven^60^ v0.12.0 with the KEGG database. Both global metabolomic profiling and stable isotope tracing data (15N serine and 15N4 hypoxanthine) were processed in this manner. MetaboAnalyst 5.0 was used for statistical analysis (PLS-DA, Heatmaps) and pair-wise comparison of sample groups^61^.

### ROS measurements by flow cytometry

ROS levels were measured by flow cytometry using the manufacturer guidelines for CellROX orange and MitoSOX Red.

### Immunohistochemistry (IHC) on DIPG patient and PDX brain sections

IHC on the mouse DIPG-PDX model was done as previously described^63^. Mice were anesthetized, perfused intracardially with PBS and 4% PFA, tumors were dissected and processed for paraffin embedding and sectioning.

For DIPG patients, tissue sections for immunostaining were obtained from deidentified paraffin-embedded brain tissues archived in the biobank repository of Cincinnati Children’s Hospital Medical Center (CCHMC). All samples were obtained after informed consent from patients’ parents/legal guardians and the Discover Together Biobank is accredited by the College of American Pathologists (CAP).

### Western Blotting

Cells were lysed in RIPA buffer containing protease and phosphatase inhibitor, protein quantitation performed by BCA. ∼ 10 ug of protein was electrophoresed on a 10 or 14% gel and transferred to a PVDF or nitrocellulose membrane. Membranes were blocked with 5% skim milk in TBST. Ponceau staining and imaging was performed as needed. Membranes were incubated with primary antibodies in 5% BSA in TBST at 4°C overnight and washed with TBST three times, incubated with secondary antibodies conjugated to horseradish peroxidase (HRP) for 2 hours at room temperature, and washed three times with TBST. Images acquired using a chemiluminescent substrate and Azure c500 Gel Imaging system.

### qRT-PCR

1 microgram of RNA (RNeasy kit , Qiagen) was used for cDNA synthesis with oligo-dT primers and Multiscribe Reverse Transcriptase (Applied Biosystems). qRT-PCR was done using SYBR® Green PCR Master Mix (Applied Biosystems) in a QuantStudio 6 Flex System (Applied Biosystems). Relative mRNA expression was calculated using the comparative Ct method after normalization to a housekeeping control. Samples were run in triplicates with a primer-limited probe for the reference gene (b-actin). Primers are provided in Supplementary table 3.

### Chromatin accessibility (ATAC-seq) in NSCs expressing H3WT and H3K27M

Omni-ATAC-Seq was performed in replicates as previously published^64^. Approximately, 50,000 cells were transferred to a microfuge tube and washed once with PBS. Cell pellets were resuspended in 48.5 μL of ice-cold resuspension buffer (10mM Tris-HCl, pH 7.5, 10 mM Nacl, 3mM MgCl2 plus 0.5 μL 10% NP-40, 0.5 μL 10% Tween-20 and 0.5 μL 1% Digitonin) and incubated on ice for 3 minutes. 990 μL of ice-cold resuspension buffer plus 10 μL10% Tween-20, invert tubes gently 3 times, followed by centrifugation at 500g for 10 minutes at 4° C. For transposition reaction, nuclei were resuspended in 50 μL of Nextera transposition reaction mix consisting of 25 μL 2x TD Buffer, 2.5 μL Nextera Tn5 transposase (Illumina # FC-121-1030), 16.5 μL 1X PBS, 0.5 μL of 10% Tween-20, 0.5 μL of 1% Digitonin and 5 μL of nuclease free water. The reaction mixture was mixed 6 times by gentle pipetting and incubated at 37° C for 30 minutes on a thermomixer at 1000 rpm. The transposed DNA was then purified by Qiagen MinElute kit (Qiagen # 28004) and eluted in 10μL of elution buffer. Transposed DNA was amplified using PCR according to the protocol’s recommendations and libraries were purified with AMPure XP beads. Libraries were sequenced on the NovaSeq 6000 (paired-end, 150 bp read length) at the CCHMC DNA Genomics Sequencing Facility, Cincinnati, Ohio. ATAC-seq data were processed and aligned to the hg19 genome using the ENCODE ATAC-seq pipeline (V2.2.0)^65,66^. Peaks were called within the pipeline using default parameters of MACS2 (q < 0.01) ^67^. Differential chromatin accessibility analysis was performed using Diffbind 3.0.15^68^ in R 4.0.2^69^. Peaks were considered differentially accessible if the FDR was less than 0.01 and the fold change was 2 or more. A modified version of HOMER^70^ using a log base 2 likelihood scoring system was used to calculate motif enrichment statistics for a large library of human position weight matrix (PWM) binding site models contained in build 2.0 of the CisBP database^71^. Data were visualized using the UCSC Genome Browser^72^.

### NFIC Chromatin immunoprecipitation sequencing (ChIP-seq) in NSCs expressing H3K27M

NFIC ChIP-seq was performed in duplicate using standard experimental procedures as previously described^73^, with a few modifications. Briefly, 2 x 107 cells were crosslinked and lysed. Nuclei were resuspended in sonication buffer (10 mM Tris [pH 8.0], 100 mM NaCl, 1mM EDTA, 0.5mM EGTA, 0.1% NaDOC and 0.5% N-lauroylsarcosine. An S220 focused ultrasonicator (COVARIS, Woburn, MA) was used to shear chromatin (150–500-bp fragments) with 10% duty cycle, 175 peak power, and 200 bursts per cycle for 7 min.

Chromatin was pre-cleared with Protein A Dynabeads for 45 minutes at 4°C on a rotator. Chromatin-immunoprecipitations were performed in an SX-8X IP-Star Compact automation system (Diagenode) with 200 μL of pre-cleared chromatin (approximately 3 million cells), 21 μL of Protein A Dynabeads (Thermo Fisher Scientific) and 5 μg of antibody. The following antibody was used for ChIP: NFIC (Bethyl Laboratories Catalog # A303-124A). Immunoprecipitation of chromatin was carried-out for 8 hours, following which, the Dynabeads were sequentially washed with Wash Buffer 1 (10 mM Tris-HCl, pH 8.0, 150 mM NaCl, 1 mM EDTA, 0.1% SDS, 0.1% sodium deoxycholate, and 1% Triton X-100), Wash Buffer 2 (10 mM Tris-HCl, pH 8.0, 250 mM NaCl, 1 mM EDTA, 0.1% SDS, 0.1% sodium deoxycholate, and 1% Triton X-100), Wash Buffer 3 for replicate 1 (1x TE Buffer, 250 mM LiCl, 0.5% NaDOC and 0.5% NP40) and for replicate 2 (50 mM Tris-HCl, pH 8.0, 2 mM EDTA, and 0.2% N-Lauroylsarcosine sodium salt), and Wash Buffer 4 (TE + 0.2% Triton X-100) for 5 min each. After the immunoprecipitation, ChIPmentation was performed, the libraries were prepared identically as the ATAC-seq method. Libraries were sequenced on the NovaSeq 6000 (single-end, 100 bp read length) at the CCHMC DNA Genomics Sequencing Facility, Cincinnati, Ohio.

ChIP-seq data were processed and aligned to the hg19 genome using the ENCODE ChIP-seq pipeline (V2.0.0)^65,66^. Peaks were called within the pipeline using default parameters of MACS2 (q < 0.01) ^67^. A modified version of HOMER^70^ using a log base 2 likelihood scoring system was used to calculate motif enrichment statistics for a large library of human position weight matrix (PWM) binding site models contained in build 2.0 of the CisBP database^71^. Data were visualized using the UCSC Genome Browser^72^.

### ATIC and PNP promoter assay

Promoter cloning and Nanoluc luciferase reporter construct generation:

Genomic DNA was amplified with the specific primers (Supplemental table S8) containing Nhel and Kpnl restriction sites. A ∼700bp sequence containing either ATIC or PNP human promoter was cloned into a pNL1.1 vector (Promega Corp., Madison, WI, USA) creating the plasmids pNL1.1-ATIC and pNL1.1-PNP, respectively. All plasmids were fully sequenced for verification purposes.

Luciferase reporter assay

2 million cells were transfected with an empty vector reporter, GFP reporter, or the ATIC or PNP reporter plasmids via electroporation using the Neon Transfection System Kit (Invitrogen, Waltham, MA, USA). Cells were plated in a 96-well white-walled plate (Corning, Corning, NY, USA) at 10,000 cells/well. All reporter plasmids were used in equimolar amounts. 48hrs after plating, luciferase activity was analyzed using NanoGlo Dual-Luciferase Reporter Assay System (Promega, Madison, WI, USA), and was measured using a GloMax Discover plate reader (Promega, Madison, WI, USA). Nanoluc luciferase activity was normalized relative to firefly luciferase activity and expressed as fold change relative to pNL1.1 empty vector.

### Cpd-14 PK Studies

Adult male (201-250 g) jugular vein cannulated Sprague-Dawley rats (Charles River Laboratories) were used for this study. The experiments were conducted in strict accordance with the Institutional Animal Care and Use Committee (IACUC)-approved protocols of University of Cincinnati.

For microdialysis, concentric-style probes (210 μm O.D. and 4.5 mm active length) were constructed with hollow fiber membrane with 13000 Da molecular weight cut off (Spectrum Laboratories, Rancho Dominguez, CA). The inlet tubing was PE-20 (0.38 mm I.D., 1.09 mm O.D., 8 cm long, Becton Dickinson, Sparks, MD) and the outlet tubing was fused silica (75 μm I.D., 147 μm O.D., 5 cm long, Polymicro Technologies, Phoenix) within Tygon tubing (21cm long, Fischer Scientific). All the probe components were glued to 26G hypodermic tubing (0.01in I.D., 0.018in O.D., 19mm long) using epoxy. Briefly, JVC rats (N=6) were implanted with stainless steel guide cannula under ketamine/xylazine (70/6 mg/kg i.p.) anesthesia 48 to 72 h before the microdialysis experiment. On the day of the experiment, a concentric-style dialysis probe was inserted through the guide cannula into the striatum.Buprenorphine (Buprenex®) was administered for perioperative analgesia. The coordinates for the tip of the probe were 1.2 mm anteroposterior and 3.1 mm lateral from bregma. Following implantation, the probes were continuously perfused with Dulbecco’s phosphate buffered saline containing 1.2 mM CaCl2 and 5 mM glucose at a flow rate of 1 μl/min overnight. On the day of the experiment, the flow rate was increased to 2 µl/min and the probes were allowed to equilibrate for an additional 2 hours before drug administration.

For drug administration and sample collection, cpd-14 was dissolved in water with 0.9% saline, administered at a final dose of 50 mg/kg via IP injection. Blood and microdialysate samples were collected simultaneously for pharmacokinetic assessment at pre-dose, 0.5, 0.75, 1, 2, 3, 4, 6, and 8h post administration followed by immediate centrifugation and separation of plasma from the blood samples. All the samples were stored at −80° C until analyzed.

For sample analysis, cpd-14 was extracted from the plasma and dialysis samples using 4:1 ratio of ice-cold methanol to sample v/v. Calibration standards were prepared by spiking varying concentrations of cpd-14 in blank plasma or dialysate buffer. Following centrifugation (10,000 rpm for 10 minutes at 4°C), the supernatant was collected and analyzed using a validated LC/MS method. All the concentrations were time-averaged over the collection interval and were analyzed using a non-compartmental approach employing Phoenix® WinNonlin version 8.3 (Certara). Key pharmacokinetic parameters estimated were the peak concentration (Cmax), time to reach Cmax (tmax), and the elimination half-life (t1/2). Results were expressed as mean ± standard deviation.

### Lentivirus Preparation and Production of Stable shRNA-Expressing Cell Lines

Lentivirus for shRNAs, CRISPR and recombinant gene expression were prepared in 293T cells as previously described^63^. 5 shRNA sequences were screened for each of the DNPB target genes. All shRNA clones (in pLKO.1 plasmid) were from the Sigma Mission RNAi shRNA library. pLKO.1-puro scrambled (NT) shRNA (Sigma) was used as a negative control. Efficacy of knockdown/overexpression was assayed by WB or qRT-PCR in target cell lines. The sequences that exhibited maximal knockdown/overexpression were used viability assays. gRNAs targeting ATIC were designed using GeneScript’s GenCRISPR gRNA Design Tool and cloned into TLCV2 (Plasmid #87360) and screened for knockout efficiency. pLenti6/V5-p53_R249S was a gift from Bernard Futscher (Addgene plasmid # 22935 ; http://n2t.net/addgene:22935 ; RRID:Addgene_22935)^74^

### Cell viability and proliferation assays

CellTiter Flour reagent was used for all viability and proliferation assays. 5000 (4- or 5-day assay) or 10000 (2 or 3 day assay) cells were plated in a 96-well plate in 150 µL media for indicated treatment, performed in triplicate.

### In vivo genetic and pharmacological treatments

Experiments using genetic knockdown/knockout of ATIC in JHH-DIPG-1, and cpd-14 treatment, were performed in our DIPG mouse model described above. The variability in detection of tumors by BLI for calling a baseline and twice weekly imaging became a tedious constraint; therefore, LSN experiments were later performed on a more reproducible model for standardized treatment schedules. For orthotopic implantation, 500,000 JHH-DIPG-1 cells were stereotactically injected into the pons of NSG mice. Mice were treated with 0.05 mg/kg buprenorphine and anesthetized with 2% to 3% isoflurane. The skull of the mouse was exposed through a small skin incision, and a small burr hole was made using a 25-gauge needle. Cells injected into coordinates 1 mm posterior and 1 mm lateral to lambda, and 5 mm deep.

Mice implanted with DIPGs stably expressing shRNA targeting ATIC were only monitored for survival analysis. For inducible ATIC knockout studies and cpd-14 treatment, mice were imaged twice weekly until tumors were detected via IVIS before indicated treatment. For LSN experiments, tumors were consistently detected at day 40 in JHH-DIPG-1 NSG model.

Quantitation of BLI for in vivo experiments was performed using Aura Imaging Software with the Living Image IVIS files. A 2 cm x 2 cm square ROI surrounding the head/tumor was used for measuring total emission (photons/s) with background subtraction.

### Differential gene expression and ATIC dependency correlation analysis

RNA-sequencing data for the parental JHH-DIPG-1 cell line and its ATIC inhibitor–resistant derivative were analyzed using the DESeq2 pipeline. Genes with an adjusted p-value < 0.05 were considered significantly differentially expressed.

For the Childhood Cancer Model Atlas (CCMA) dataset, gene expression values (CPM+1) and CRISPR gene dependency z-scores were downloaded from the publicly available CCMA portal^52^. Spearman correlation analysis was conducted to assess the relationship between gene expression and ATIC dependency scores across all cell lines. Genes with a correlation p-value < 0.05 were retained for further analysis.

Genes that were significantly differentially expressed in the ATIC-resistant vs. parental DIPG line and also significantly correlated with ATIC dependency in the CCMA dataset were identified by overlapping the two filtered gene sets. Overlap directionality was confirmed based on the direction of fold-change and correlation coefficient.

### Statistical Analysis

Data was analyzed using T test or ANNOVA. Details of statistical analyses can be found in the figure legends. Statistical analysis and enrichment analysis of in vitro untargeted metabolomics was performed in MetaboAnalyst 5.0 and assigned to KEGG curated pathways. For in vivo tumor metabolomics, MetaboAnalyst 5.0 was used for statistical analysis (PLS-DA, Heatmaps) and pair-wise comparison of sample groups.

## Notes

### Competing Interest Statement

The authors have declared no competing interest.

