## Supplementary figures and images for "Integrative Multi-Omics Analysis Identifies Nuclear Factor I as a Key Driver of Dysregulated Purine Metabolism in DIPG"

### Supplemental figure 1.tif

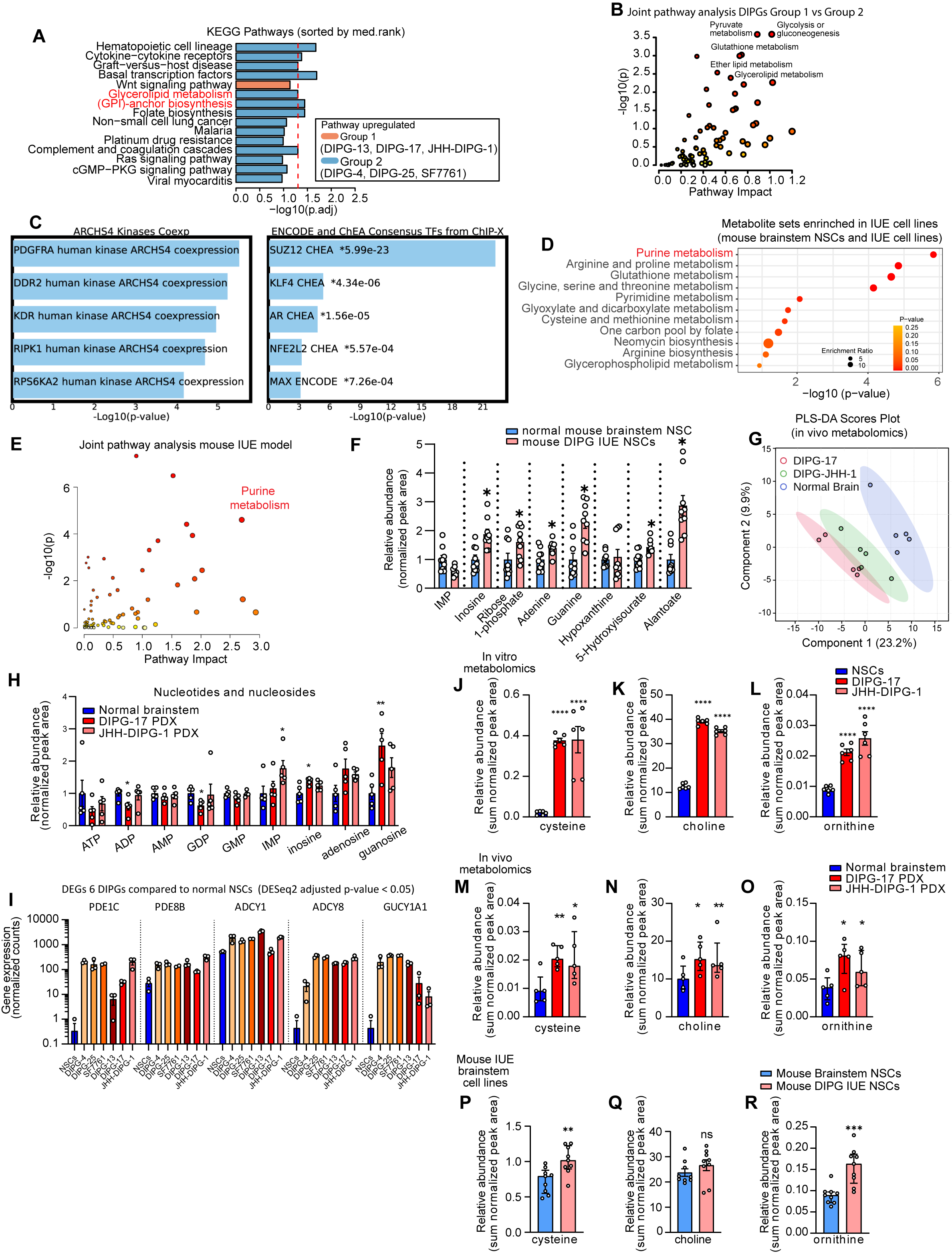

### Supplemental figure 2.tif

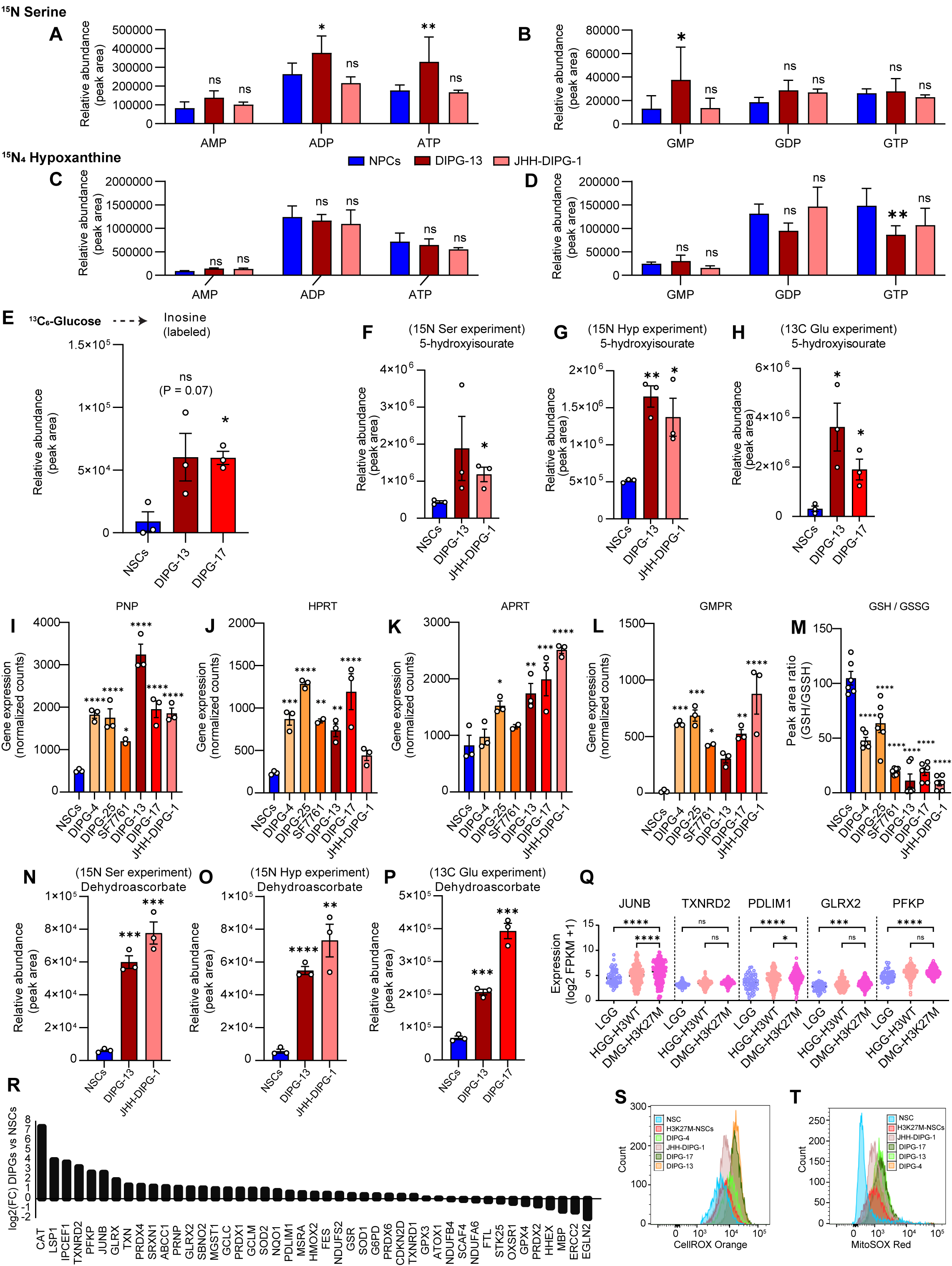

### Supplemental Figure 3.tif

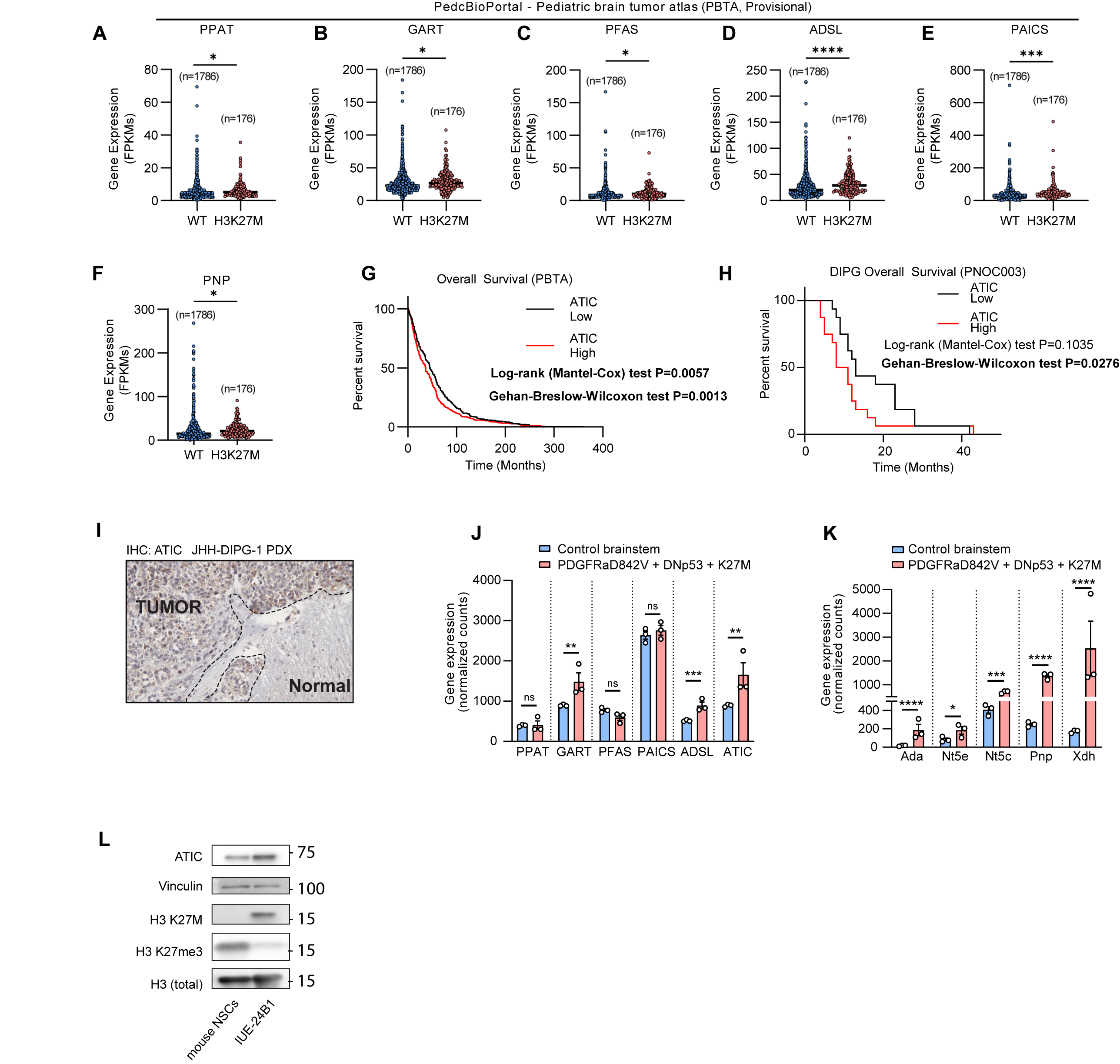

### Supplemental figure 4.tif

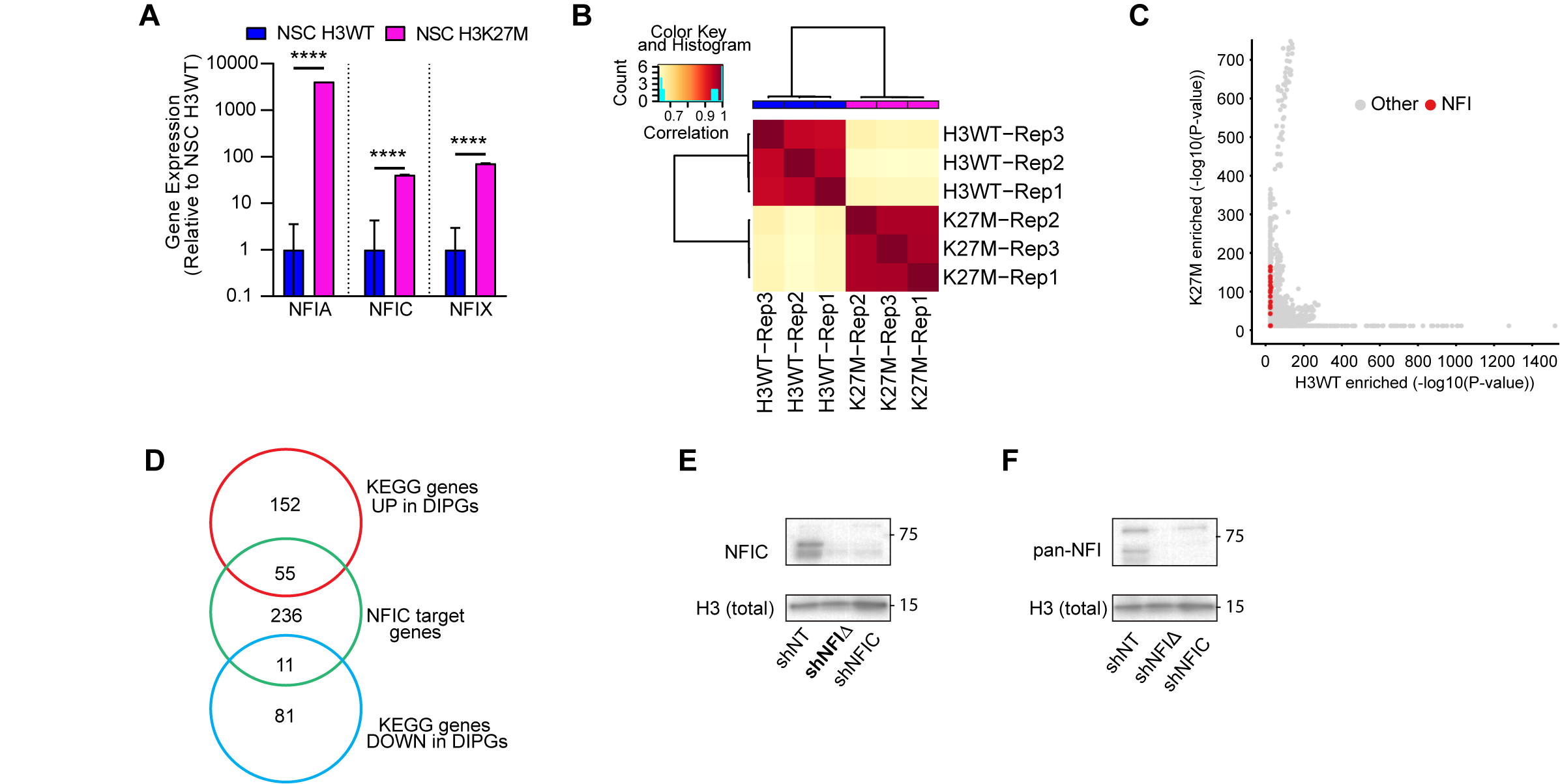

### Supplemental figure 5.tif

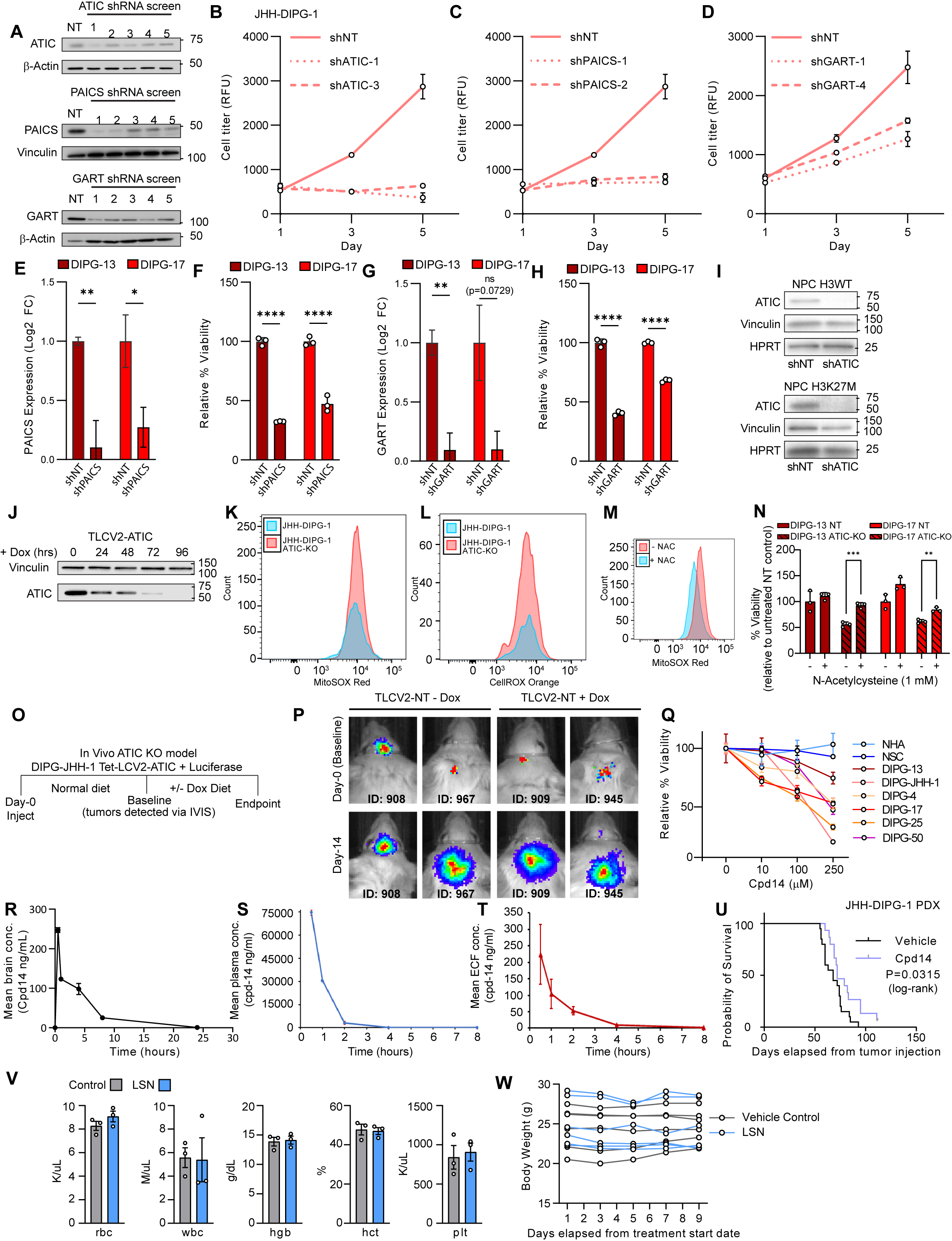

### Supplemental figure 6.tif

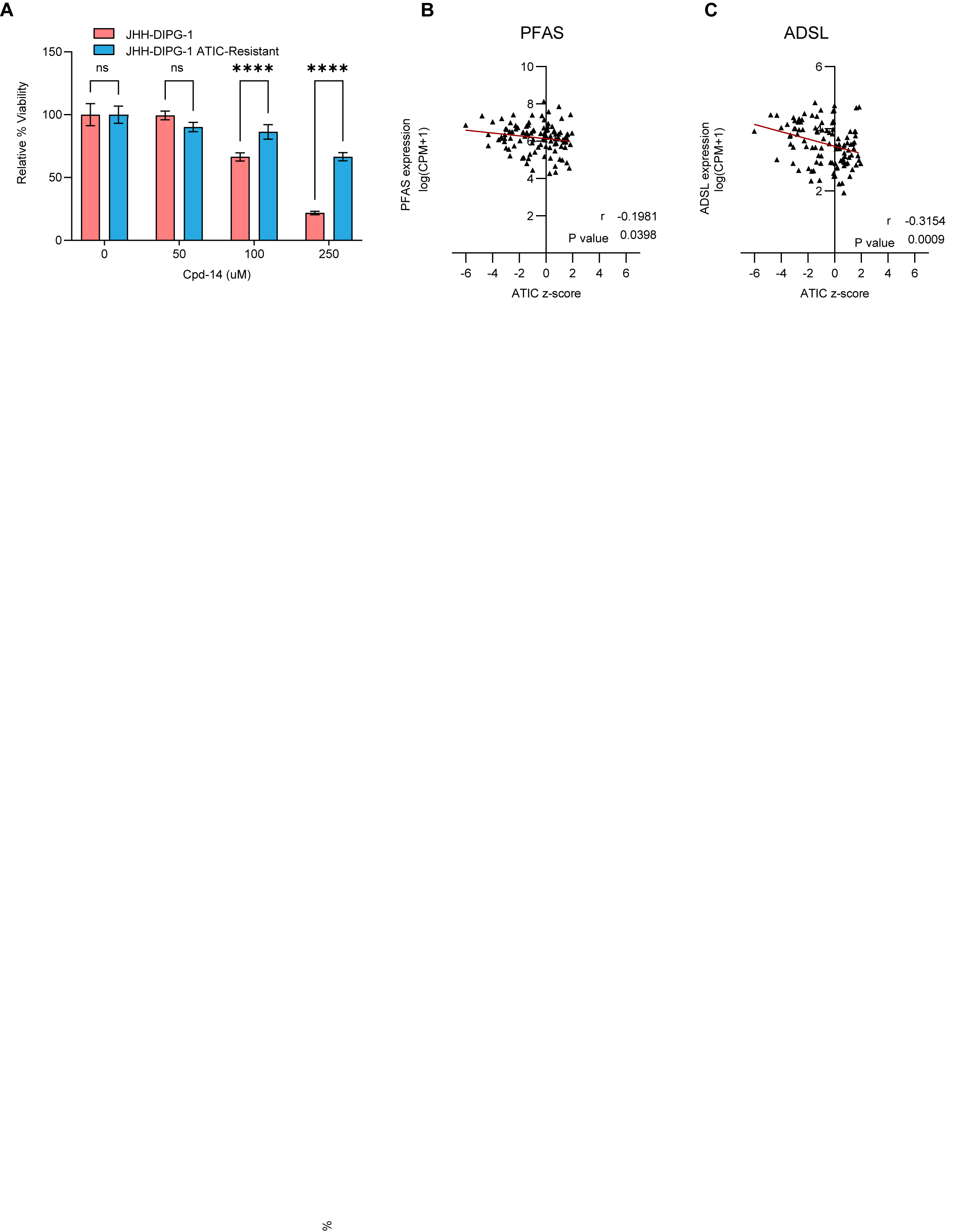
